# When species trees disagree: an approach consistent with the coalescent that quantifies phylogenomic support for contentious relationships

**DOI:** 10.1101/2020.03.27.012237

**Authors:** Richard G.J. Hodel, Joseph F. Walker, L. Lacey Knowles, Stephen A. Smith

## Abstract

Phylogenies inferred using both concatenation- and coalescent-based analyses typically render highly congruent trees. However, when they disagree, they often differ with respect to historically contentious and evolutionarily important relationships. These relationships may also involve etiolated lineages where increased sampling is not possible. Recently, methods aimed at interrogating single relationships or trees have emerged as promising investigative tools to examine these cases. Although recent methods such as “Edge-based Phylogenomic Support analYsis” (EPSY) led to insights into both systematic error and real biological signal, whether they are consistent with the coalescent in cases with high Incomplete Lineage Sorting (ILS) has yet to be characterized. Here, we use simulations and an empirical dataset to test the performance of EPSY, concatenation, and coalescent-based summary analyses under high levels of ILS. We focused on high-ILS scenarios because these represent the typical difficult cases that researchers often face due to the prevalence of ILS in phylogenomic datasets. ILS is known to be a major cause of phylogenomic conflict, which confounds many biological conclusions that depend on a resolved phylogeny, such as inferring ancestral character states, biogeographic reconstructions, and domestication histories. Our study found that EPSY was consistent with the coalescent in a high-ILS empirical dataset. In high-ILS simulations EPSY infers the correct edge more than half the time, whereas coalescent based methods and concatenation methods inferred the actual tree 37.8% and 25% of the time, respectively. All methods have conditions under which they generate the most accurate inferences. Given the levels of ILS in simulations, 26.2% of the time no method recovered the true tree. This zone where no current method can infer the true topology is likely due to properties of the species tree, such as the length of internal edges adjacent to a conflict and/or the length of the shortest branch. Nevertheless, the EPSY approach proves to be a valuable complement to phylogenomic analyses for interrogating regions of the tree with conflicting hypotheses generated from past studies or alternative inference methods. Our analyses highlight that robust phylogenetic trees may not be possible under some scenarios regardless of method and data source.

## INTRODUCTION

The rapid increase in the size of genomic datasets for phylogenetics has failed to lead to the undisputed resolution of many contentious relationships in the Tree of Life (e.g., Near 2009, Janvier 2010, Wickett et al. 2014, Xi et al. 2014, Arcila et al. 2017, Miyashita et al. 2019). As larger datasets are constructed, analyzed, and reanalyzed, the most contentious regions of the Tree of Life remain hotly debated in the literature (Shen et al. 2017). Phylogenetic relationships associated with key transitions, such as the evolutionary history leading to jawed vertebrates (Near 2009, Janvier 2010, Miyashita et al. 2019) and a key position for inferring the order of early angiosperm divergences (Wickett et al. 2014, Xi et al. 2014) remain recalcitrant. Despite methods that model nucleotide evolution and the coalescence of genealogies, discordance within and among studies is prevalent. A resolved phylogeny is a critical first step for making biologically-meaningful inferences including determining ancestral characters (Xi et al. 2014), inferring biogeographic reconstructions (DaCosta et al. 2019), resolving rapid radiations (Whitfield & Lockhart 2007), identifying domestication events (Kates et al. 2017), and inferring whether or not lineages arose as the result of a hybridization event (Zhang et al. 2018).

Biological processes such as incomplete lineage sorting (ILS), gene duplication and loss, hybridization, and horizontal gene transfer (HGT) cause genomes to be a composite of evolutionary histories that complicate species tree reconstructions. Furthermore, systematic error presents a challenge for inferring the phylogenetic signals that underlie a single dataset, prior to reconstructing the species relationships. Identifying and accounting for these processes in empirical datasets remains conceptually and analytically challenging, particularly for processes other than ILS (Salichos et al. 2014, Smith et al. 2015, Kobert et al. 2016, Knowles et al. 2018). Although coalescent-based summary analyses are typically vigorous to ILS and computationally tractable (Mirarab et al. 2014b), analyses accounting for other sources of conflict represent intense computational challenges (Ané et al. 2006, Boussau et al. 2013). Recent methods have worked to isolate a single edge and infer what properties of the disparate influence among the data (Castoe 2009, Brown & Thomson 2017, Shen et al. 2017, Walker et al. 2018). This has led to the discovery that many contentious nodes are sensitive to a single gene (Shen et al. 2017) and these genes may be the result of systematic (e.g., orthology inference) error (Brown & Thomson 2017) or possibly arise for unknown biological reasons (Walker et al. 2018). As a result, methods that allow conflict to exist outside of the edge of interest have been explored (Arcila et al. 2017, Walker et al. 2018, Smith et al. 2019).

The age of whole genomes marks the beginning of an era where accumulating more molecular data to understand a species relationship is no longer possible. While we can push the limits of our current methods with these datasets, it is becoming apparent that novel approaches for interrogating genomic data are needed. Recently, methods aimed at interrogating single relationships (edge-based likelihood comparisons (hereafter referred to as Edge-based Phylogenomic Support analYsis; abbreviated ‘EPSY’); Walker et al. 2018) or trees (comparison of gene-wise log-likelihood scores (ΔGLS) between trees; Shen et al. 2017) have shown great promise in accommodating heterogeneity across a tree and have emerged as promising investigative tools to help examine these cases. Theoretically, an edge that captures the species relationship will have a longer branch subtending the relationship of interest, penalizing the likelihood of those edges that conflict with the true tree when comparing the likelihoods of topologies that only differ at that edge. EPSY leverages these differences to analyze gene tree conflict in more detail, and therefore can quantify phylogenomic support. Because it is a data-centric approach, EPSY does not identify the process that generated heterogeneity within a phylogeny. While these analyses have proved useful in interrogating relationships (Walker et al. 2018) and identifying problematic gene regions (Smith et al. 2019), one key unanswered question is whether EPSY can disentangle the signal that generates conflict at contentious edges when the source of conflict is known. ILS is known to be prevalent in phylogenomic studies (Edwards 2009) and is a major cause of conflict throughout the Tree of Life (Smith et al. 2015); any phylogenomic method that accommodates conflict inconsistent with ILS will have limited utility. Because ILS can be accounted for in coalescent-based summary species tree methods (e.g., ASTRAL; Mirarab et al. 2014b), it presents an opportunity to test if EPSY methods produce similar results to coalescent species tree analyses when ILS is simulated to be the cause of conflict. For simplicity, coalescent-based species tree analyses in this paper are conducted in ASTRAL due to its speed, ease of implementation, and popularity in the phylogenomics community despite the fact that ASTRAL is not a true coalescent species tree method. ASTRAL can accommodate large datasets because it is statistically consistent with the coalescent; unrooted quartets do not exist in the anomaly zone (Degnan 2013) so as the number of gene trees increases, the probability of correctly inferring the species tree increases (Mirarab et al. 2014b). Hereafter, when we use ‘coalescent’ analyses, we are referring to ASTRAL analyses.

In this study, we investigate what course of action researchers should take when species tree reconciliations differ. We first study whether EPSY can infer the correct species tree phylogeny in cases when coalescent and concatenation topologies disagree. We use simulated and empirical datasets to investigate whether three approaches – EPSY, methods statistically consistent with the coalescent (ASTRAL), and methods using a supermatrix concatenation approach (unpartitioned RAxML) – can recover the true species tree topology and assess how these perform under varying tree shapes and levels of ILS. Specifically, we simulate sequence data for gene trees in species trees under conditions with ILS-generated conflict to investigate how EPSY methods behave in multiple regions of phylogenetic tree parameter space. If the process causing conflict is known (i.e., ILS), we assess if EPSY is consistent with the coalescent. Additionally, we use an empirical dataset – a multilocus data of birds focused on phylogenetic relationships of Neoaves with documented high ILS at specific contentious nodes (Jarvis et al. 2014, Prum et al. 2015, Suh et al. 2015, Reddy et al. 2017) – to compare to the simulated datasets. Finally, we assess how these results can be applied to empirical datasets and offer recommendations for researchers addressing phylogenomic conflict.

## MATERIALS AND METHODS

### Simulations

#### Species Trees

Simulations were designed to investigate the conditions under which coalescent, concatenation, and EPSY methods recover a known species tree topology. We simulated eight datasets with gene tree-species tree conflict caused by ILS that varied in several key parameters – specifically, root height, species birth rate, and effective population size – to maximize the extent of phylogenetic tree parameter space represented by the simulated datasets (Table 1, Supplemental Fig. S1). We simulated 50 species trees under eight different tree topologies (simulated datasets A-H) for a total of 400 species trees, each with 11 taxa (focal species and an outgroup; Fig. 1). Although all datasets were simulated under ILS, the datasets differ in the amount of ILS as a function of the different parameter settings (see Table 1 and Supplemental Information for the details of the parameterization of each of the eight datasets).

**Table 1.** For each of the simulated datasets (A-H), the specific parameters for root height, species birth rate, and effective population size used for the species trees. In general, shallower root height, a lower species birth rate, and a higher effective population size are expected to lead to higher ILS.

| Dataset | Root height | Species birth rate | Effective population size |
| --- | --- | --- | --- |
| A | 1.0E+08 | 1.0E-08 | 1.0E+06 |
| B | 1.0E+08 | 1.0E-07 | 1.0E+06 |
| C | 1.0E+08 | 1.0E-08 | 1.0E+07 |
| D | 1.0E+08 | 1.0E-07 | 1.0E+07 |
| E | 2.0E+08 | 1.0E-08 | 1.0E+06 |
| F | 2.0E+08 | 1.0E-07 | 1.0E+06 |
| G | 2.0E+08 | 1.0E-08 | 1.0E+07 |
| H | 2.0E+08 | 1.0E-07 | 1.0E+07 |

**Figure 1.**
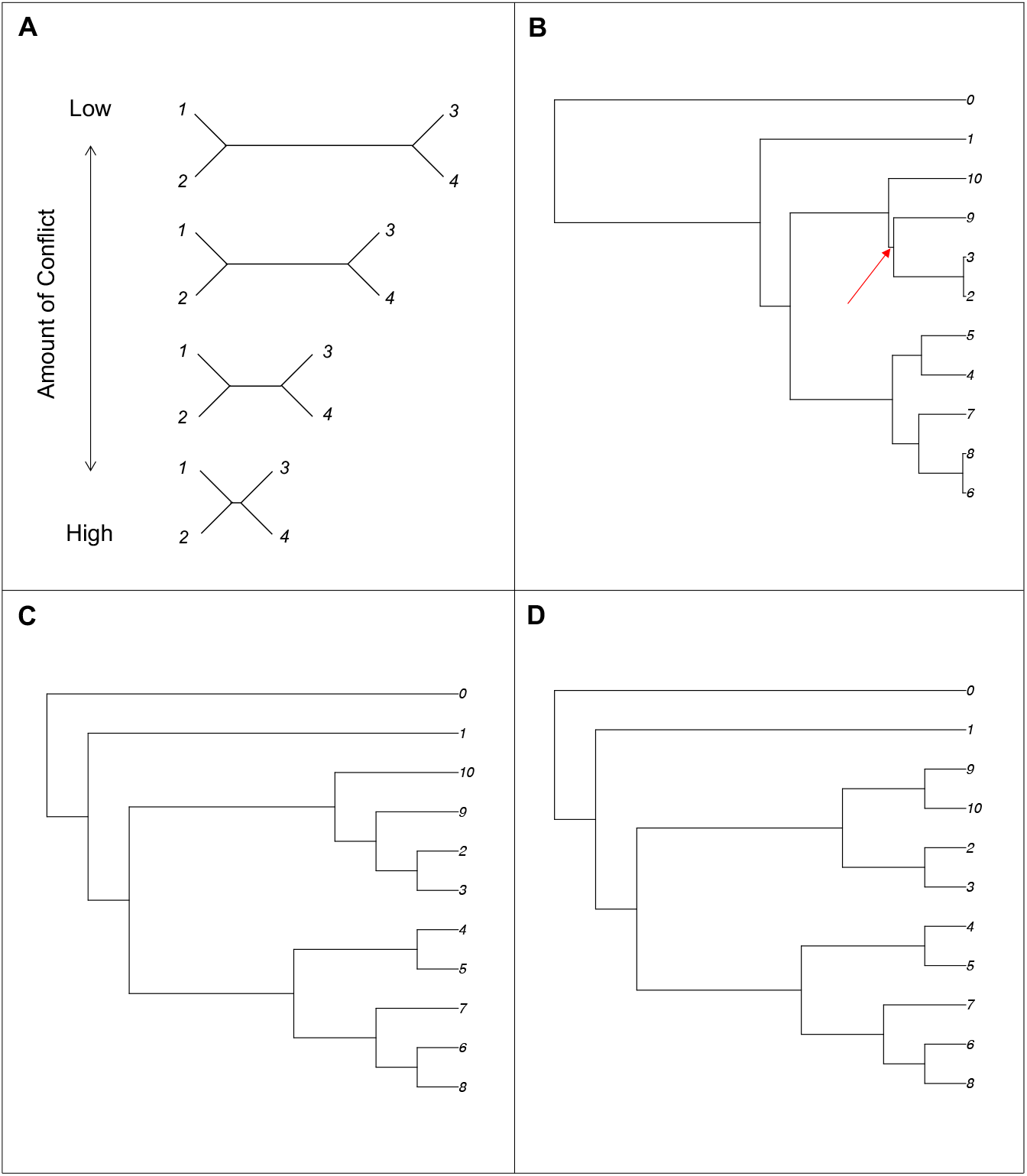
A) The theoretical expectation of how the amount of gene tree / species tree conflict should change with internal branch length. As the internal branch length decreases relative to terminal branch lengths, the amount of conflict between gene trees and the species tree increases. The four taxa trees show the range of scenarios from no conflict (long internal branch) to high conflict (very short internal branch). B) A simulated 10-taxon (plus outgroup) species tree with high levels of conflict due to ILS. The red arrow indicates the internal edge adjacent to the conflict. In this case, a supermatrix (i.e., RAxML) approach does not infer the true species tree from the simulated sequence supermatrix but the coalescent (i.e., ASTRAL) approach recovers the true species tree topology from estimated gene trees. EPSY and coalescent analysis inferred the true topology, and concatenation recovered a different topology. C) The ASTRAL-inferred phylogeny. D) The RAxML-inferred phylogeny; note the placement of taxon 9.

#### Gene Trees

For each of the 400 species trees, we simulated 100 gene trees (i.e., coalescent genealogies) for each species tree using the software package SimPhy (Mallo et al. 2016); configuration files showing all SimPhy parameter settings are available in Supplemental Information. For each gene tree, we simulated DNA sequence data using the INDELible sequence simulator under an indel-free GTR+G model of evolution (Fletcher & Yang 2009); configuration files showing all INDELible parameter settings are also available in Supplemental Information. We estimated gene tree topologies using the resulting sequence alignment in RAxML v8 with the GTRGAMMA model. The simulated DNA sequences and estimated gene trees were used in subsequent analyses. To quantify the amount of conflict, for each species tree we used RAxML (Stamatakis 2014) and the R package *phytools* (Revell 2012) to calculate the normalized Robinson-Foulds (RF; Robinson & Foulds 1981) distance between the species tree and the coalescent genealogy, and between the species tree and the estimated gene trees (note mutational variance can generate a mismatch between the actual and estimated gene trees; Huang et al. 2010).

### Coalescent, concatenation, and EPSY

For each simulated species tree, a concatenated supermatrix was constructed using the sequence data simulated with INDELible for 100 gene trees. A custom script was used to convert 100 fasta files (i.e., INDELible output for 100 gene trees) into a single supermatrix file compatible with downstream software packages (i.e., RAxML). For each species tree, the tree topology was estimated on the supermatrix using RAxML with the GTRGAMMA model (Stamatakis 2014). ASTRAL v5.6.1 (Mirarab et al. 2014b) was used to infer species tree topologies based on the estimated gene tree topologies (i.e., estimated from simulated sequence data using RAxML) in a framework statistically consistent with the coalescent.

For every simulated species tree, the EdgeTest.py script in the software package EdgeTest (https://github.com/jfwalker/EdgeTest) was used to calculate the likelihood for among conflicting hypotheses (Walker et al. 2018). Specifically, conflicting edges between the simulated tree and the concatenation and coalescent-based trees were identified using ‘pxbp’ from the phyx package (Brown et al. 2017a). Each edge was used as a constraint, and the likelihood of the constraint was calculated for each gene individually. The individual likelihoods of each gene were summed for their given edge, allowing a value for the likelihood of the edge to be calculated. EPSY, implemented in EdgeTest.py, compared the summed difference in likelihood scores for all genes and the number of genes supporting each of the alternative topologies--coalescent and concatenation. We also investigated only genes that differed by a difference of >2 lnL between alternative relationships, which is considered a cutoff for statistical significance (Edwards 1984). In some species trees, there were multiple conflicting edges in the tree; in this case, EdgeTest.py compares alternative topologies for each conflicting edge between the input trees. In almost all cases, both the sum difference in likelihood scores and the number of genes support the same relationship. However, using both metrics can be valuable for detecting cases when one or a few outlier genes are having a disproportionately large effect (Brown & Thomson 2017, Shen et al. 2017, Walker et al. 2018).

We investigated trends that emerged when simultaneously considering all simulated datasets (A-H) combined, and patterns specific to each simulated dataset. Several properties of simulated species trees that could influence if certain methods were more or less likely to perform well were investigated in detail. Specifically, for all simulated species trees with coalescent-concatenation conflict, we identified the shortest branch length (i.e., the shortest branch in the entire tree), the length of the internal edge adjacent to a conflict, and the height of a conflict (measured from the tip). We used the R packages *ape* (Paradis et al. 2004) and *phytools* (Revell 2012) to calculate the above measurements. The height of a conflict was calculated using the *ape* function ‘node.depth.edgelength’, which measures the root-to-node distance. The height of the conflict (i.e., tip-to-node-in-conflict distance) was calculated by subtracting the root-to-node distance from the root-to-tip height of the tree.

### Empirical dataset

A phylogenomic dataset with documented high ILS at certain nodes was used to test if an empirical dataset behaves similarly to our simulations. Several nodes deep in the bird phylogenetic tree (Fig. 2) have high instances of ILS (Suh et al. 2015) leading to incongruence (Reddy et al. 2017, Brown et al. 2017b). Our empirical dataset (modified from Reddy et al. 2017) consisted of 59 genes from 48 taxa that were used to infer a phylogeny of Neoaves. We used 48 taxa that represent major groups of birds featured in Jarvis et al. (2014) because it was a manageable number of taxa that could clearly show conflict between inference methods. However, we used loci from Reddy et al. (2017) for these 48 taxa to build a supermatrix to minimize missing data in order to facilitate estimation of each gene tree from empirical sequence data. Additionally, we wanted to minimize data type biases that have been documented to exist in these taxa/genes (Reddy et al. 2017).

**Figure 2.**
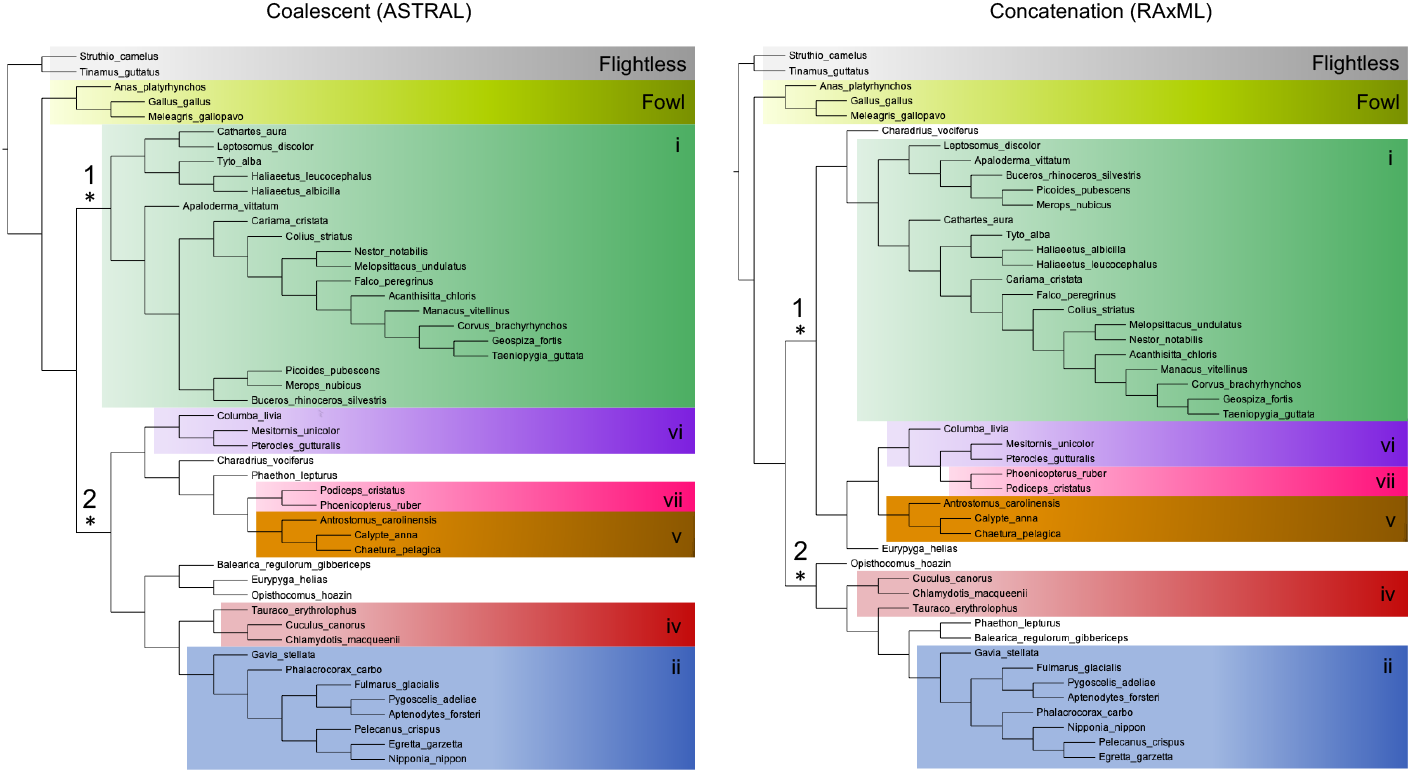
The bird phylogenies inferred using empirical data and either coalescent (ASTRAL, left) or concatenation (RAxML, right) approaches. Major clades or groups are colored and labeled (i-vii) in accordance with Reddy et al. (2017). The asterisks indicate contentious nodes along the backbone of the phylogeny where conflict is due to ILS (Suh et al. 2015).

### Coalescent, concatenation, and EPSY

The 59 genes were concatenated into a supermatrix using the ‘pxcat’ command in the phyx software package (Brown et al. 2017a; https://github.com/FePhyFoFum/phyx). Then, RAxML was utilized to infer the phylogeny using the rapid bootstrapping command with 100 bootstrap replicates. Additionally, we estimated gene trees for each of the 59 genes using RAxML with the rapid bootstrapping command with the same search strategy. Then, the 59 gene tree topologies were used as input for ASTRAL (Mirarab et al. 2014b), which was used to infer species tree topologies. We identified conflicts between the coalescent and concatenation topologies using phyparts (https://bitbucket.org/blackrim/phyparts; Smith et al. 2015) and PHAIL. In cases where conflict was detected, the ‘conducting alternative relationship analyses’ feature in phyckle (Smith et al. 2019; https://github.com/FePhyFoFum/phyckle) was used to compare the alternative topologies with EPSY; phyckle uses RAxML to calculate likelihood scores for all genes. Two metrics--the sum of the differences in likelihood scores for all genes, and the number of supporting genes--were then used to identify the preferred topology between the two competing topologies (i.e.., coalescent versus concatenated). We focused on identifying and analyzing conflicts along the backbone of the bird phylogeny, as this is where there are high levels of ILS (Fig. 2; Suh et al. 2015), but we also investigated any relationships where there was an unexpected conflict.

## RESULTS

### Cases where EPSY identifies the true topology when neither coalescence nor concatenation can

The percentage of species trees with at least one conflict ranged from 12% (dataset A) to 68% (dataset G; Supplemental Fig. S2) and was consistent with expectations based on RF distances between gene trees and species trees (Fig. 3, Supplemental Fig. S3). For some species trees, there was no disagreement between topologies of trees inferred from coalescent-based and concatenation-based analyses (hereafter, referred to simply as coalescent and concatenation topologies). In these cases, ILS was not substantial enough to generate conflict (Supplemental Fig. S2). In other cases with conflict between the two phylogenetic trees inferred by coalescent and concatenation methods, and for which one of the two methods recovered the true species tree (Fig. 4), on average EPSY identified the method that inferred the true topology 54.9% of the time. In the cases in which neither coalescent nor concatenation analyses inferred the true topology, EPSY supports the true topology for some, but not all species trees (Fig. 4; i.e., no method was able to correctly identify the true topology in some cases; see also Supplemental Fig. S4).

**Figure 3.**
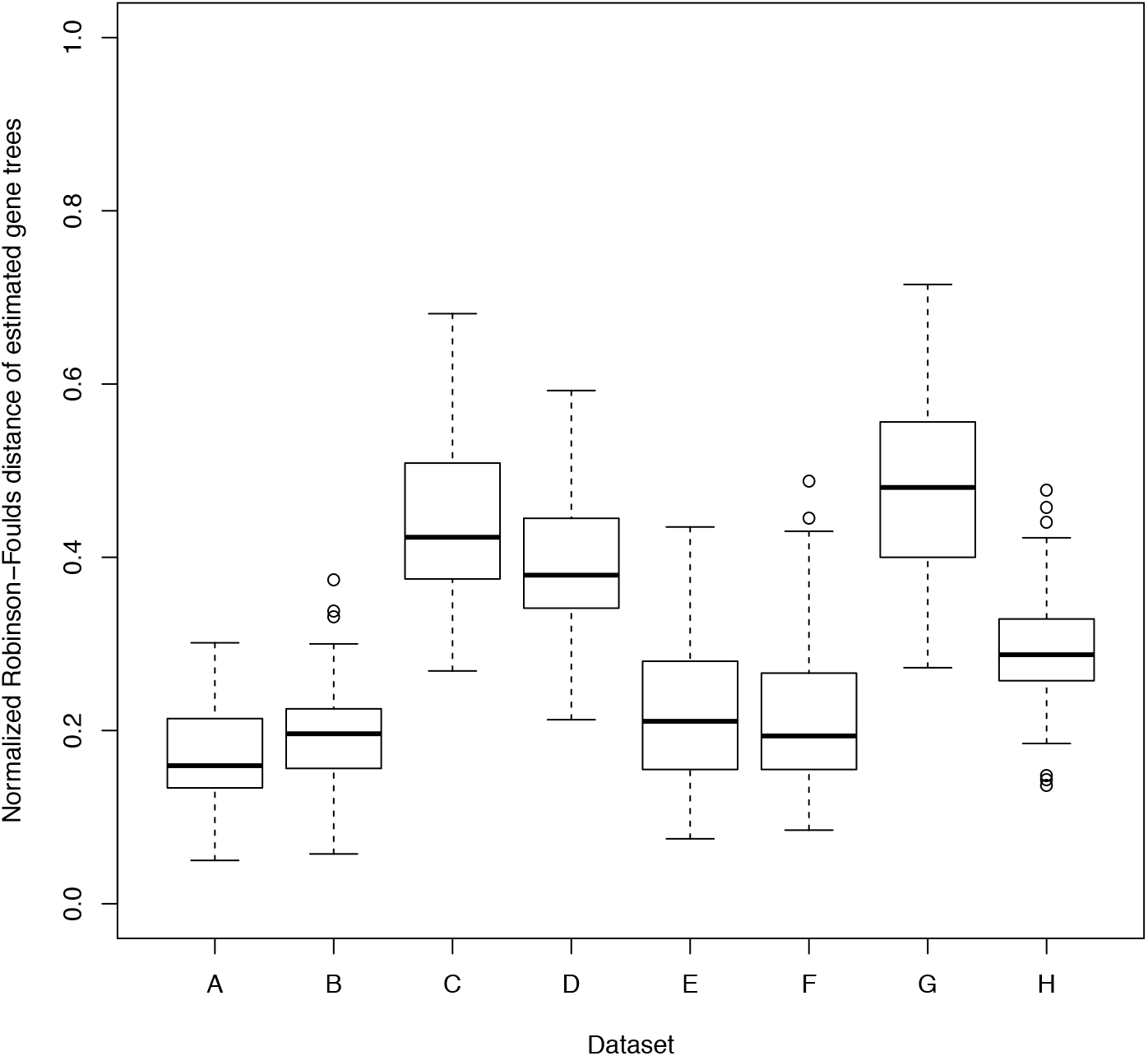
Box-and-whisker describing the normalized average Robinson-Foulds distances between each species tree and the estimated gene trees in the species tree are displayed for all eight datasets. Each box plot shows the median (thick line), and 25% and 75% quartiles (lower and upper horizontal bounds of box, respectively), and the whiskers show 1.5*(interquartile range).

**Figure 4.**
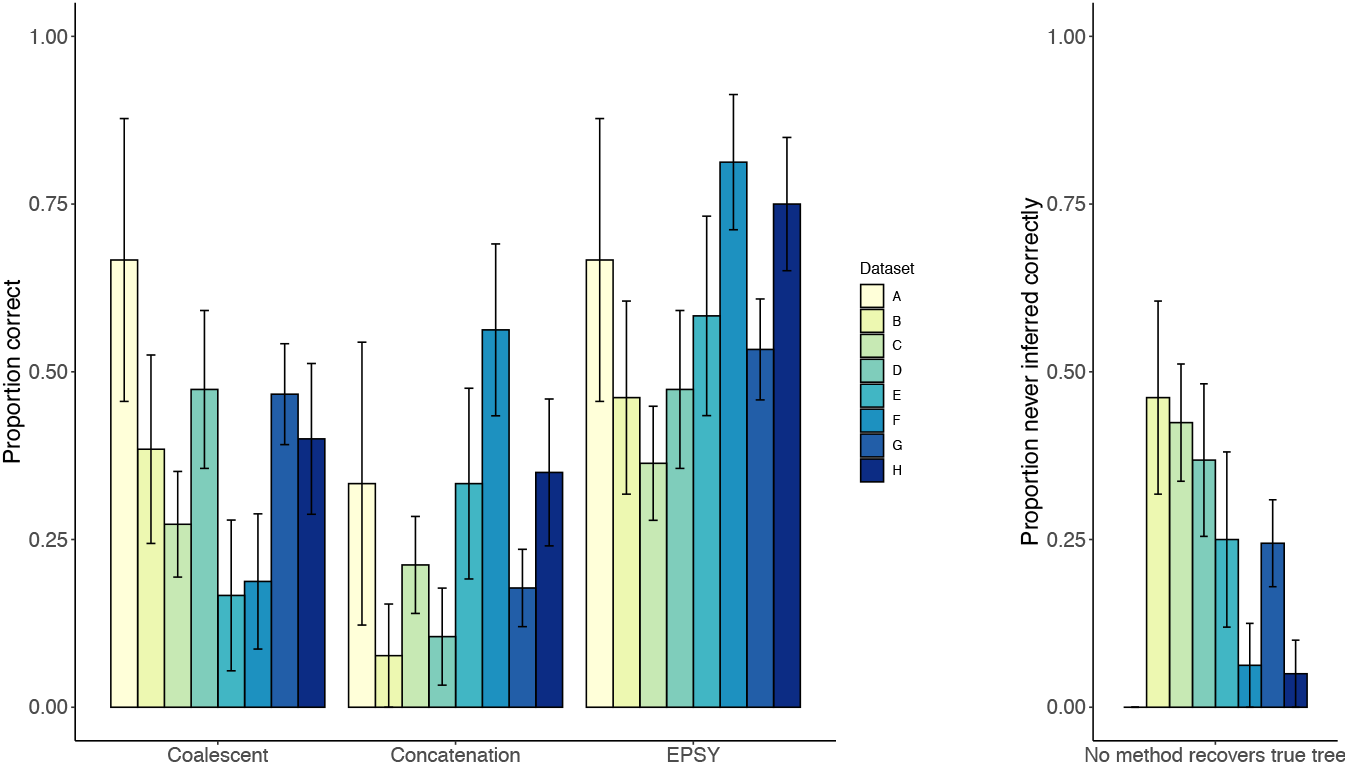
The proportion of conflicts correctly inferred using each method for all of the eight simulated datasets are indicated on the left. Cases where no method obtained the correct topology for an edge with conflict are depicted to the right of the legend, and error bars depict standard error of the mean. See Table 1 for details about the parameterization of each of the simulated datasets.

Trends emerged when we evaluated species trees with coalescent-concatenation conflict from all eight simulated datasets. Specifically, EPSY identified the correct phylogenetic relationship in the highest proportion of conflicts: EPSY recovered the true topology 54.9% of the time, compared to 37.8% for the coalescent and 25.0% for concatenation (Figure 4). In 26.2% of cases with conflict, no method could recover a true topology; we later describe properties of a tree associated with conflicting inferences, as well as tree properties associated with certain methods outperforming others.

### EPSY is consistent with the coalescent

EPSY was largely consistent with the coalescent as opposed to concatenation under high ILS conditions simulated here. When the EPSY results corresponded with either the coalescent or concatenation topology, it agreed with the coalescent topology more frequently than the concatenation topology (57.1% versus 42.9% of the time). For the four datasets with the highest ILS (C, D, G, H; based on RF distances; Fig. 3) the pattern was similar, albeit with 63.8% versus 36.2% agreeing with the coalescent versus concatenation topology. When considering all simulated datasets, the proportion of times that EPSY correctly identified the true topology was greater than the frequency at which coalescent analyses recovered the true topology (two-tailed t-test: t = 2.45, p-value = 0.028; Fig. 4, Supplemental Fig. S5) and was much greater than when the tree topology was inferred by concatenation (t = 3.86, p-value = 0.001758; Fig. 4). Additionally, when examining only datasets with the highest ILS (C, D, G, H), the relationships between the proportion of species trees in conflict correctly detected by each method indicated that EPSY methods were consistent (i.e., statistically equivalent) with coalescent methods (t = 1.21, p-value = 0.28; Fig. 4, Supplemental Fig. S5), and significantly outperformed concatenation (t = 2.98, p-value = 0.02646).

*Properties of a tree indicate when conflict will occur and when certain methods will recover the true tree*

### Length of the shortest branch in species tree

The length of the shortest branch in a species tree predicted whether or not coalescent and concatenation phylogenies would conflict, and whether any method (coalescent, concatenation, EPSY) could recover the true species tree topology (Fig. 5, Supplemental Fig. S6). In general, as the shortest branch length in the species tree decreased, a conflict between coalescent and concatenation was more likely, and it was more likely that no method could find the true species tree (Fig. 5). Furthermore, as the shortest branch length decreased, the proportion of cases where no method found the true tree relative to frequency of coalescent/concatenation conflicts increased (Fig. 5). These trends were also true when all datasets were considered simultaneously; the average shortest branch in species trees with no coalescent/concatenation conflict was much longer than the shortest branch in species trees that had conflict or incorrectly inferred species trees (Supplemental Fig. S6). Across all datasets, the conflict-free species trees had significantly longer shortest branch lengths than the species trees with conflict (t = 9.38, p-value <0.0001) and the species trees with conflict had significantly longer shortest branch lengths than the species trees where no method recovered the true tree (t = 4.24, p-value < 0.0001). The average length of the shortest branch length was approximately three times larger in species trees with no conflict (1,569,329 generations) compared to species trees with conflict (483,003 generations), which was in turn over twice as large as species trees where no method recovered the true topology (220,138 generations). In nearly all cases when datasets were evaluated individually, there was an association between the conflict-free species trees and conflicting species trees (but see dataset G; Fig. 5).

**Figure 5.**
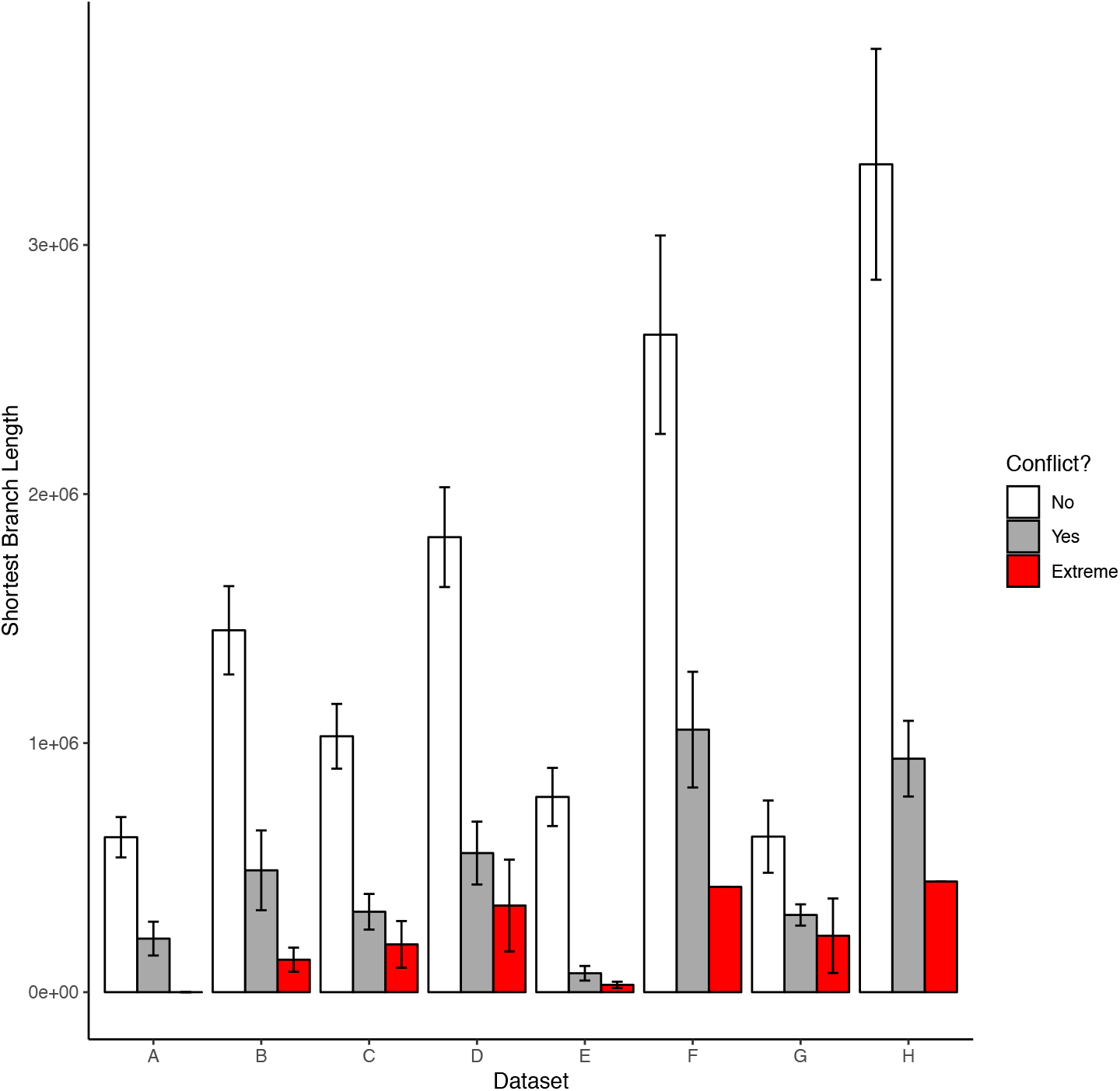
For each of the eight datasets, the mean (+/- standard error) length of the shortest branch length in a species tree for the three possible outcomes: no conflict (white), conflict (gray), or extreme conflict (i.e., no method recovers the true topology; red).

### Length of the internal edge adjacent to conflict

In a given species tree with coalescent-concatenation conflict, the length of the internal edge adjacent to conflict (i.e., closer to the root) predicted when certain methods would recover the true tree (Fig. 1B). In general, as the length of the internal edge adjacent to conflict increased, it was more likely that a coalescent or EPSY would identify the true tree than a concatenation analysis (Fig. 6).

**Figure 6.**
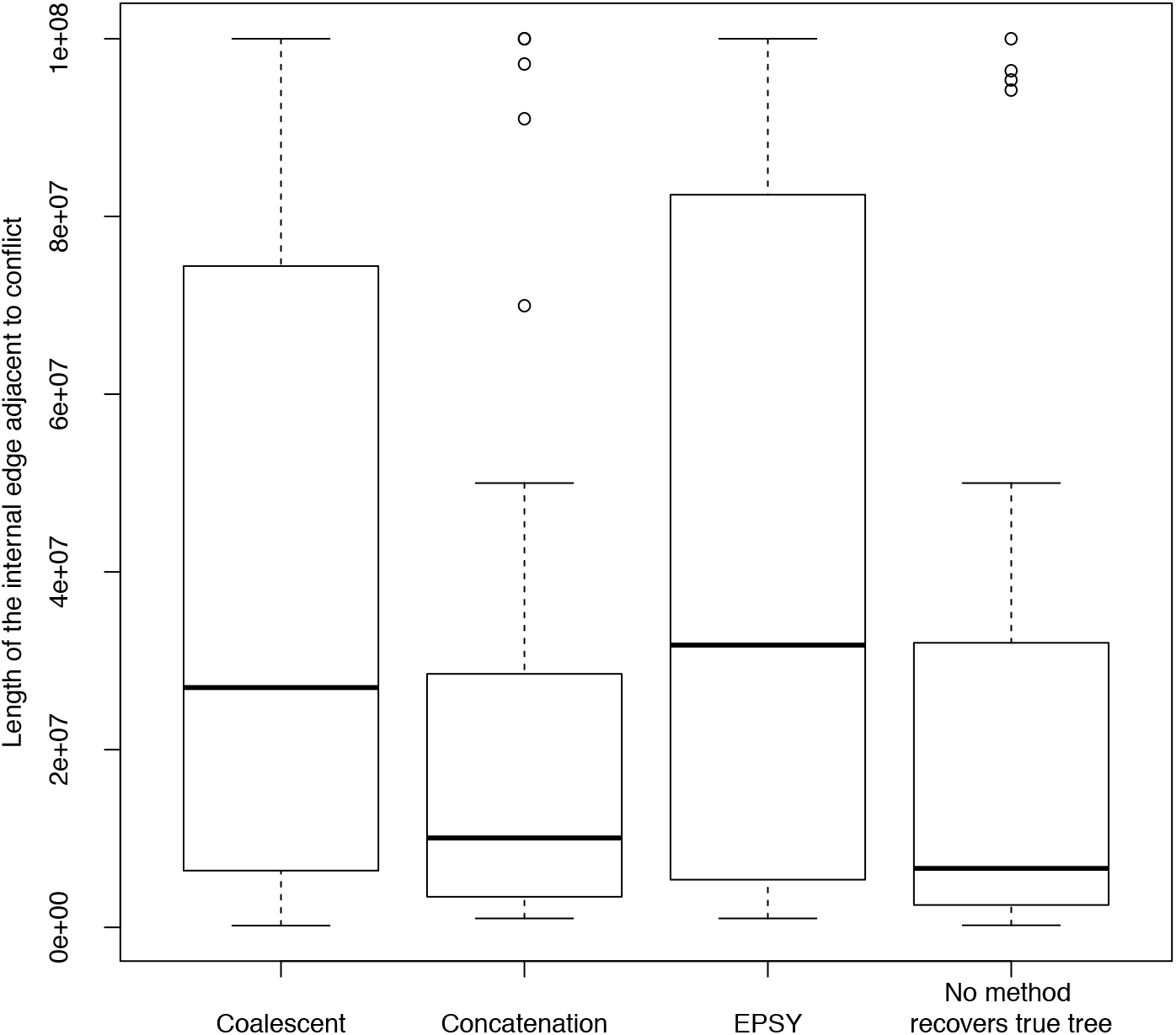
Box-and-whisker plots showing the range of the length of the internal edge adjacent to the conflict of interest. Each plot shows the length of the internal edge where the true topology was recovered by coalescent, concatenation, EPSY, or not recovered by any method, respectively.

Across all datasets in cases when the true tree was recovered, the lengths of adjacent edges in both EPSY and coalescent analyses were significantly longer than those of concatenation analyses (edge: t = 3.01, p-value = 0.00342; coalescent: t = 2.51, p-value = 0.0137). The mean length of the internal edge adjacent to conflict were similar when coalescent and EPSY methods recovered the true tree (39,674,386 and 41,329,948 generations, respectively). Notably, both of these edge lengths were nearly twice as large as the internal adjacent edge length in cases when concatenation found the true topology (22,899,746 generations) or no method found the true topology (20,968,949 generations).

### Height of conflict

The height of the edge involving conflict (i.e., the distance from the tip to the node in conflict) was also associated with the performance of different methods (Supplemental Fig. S7). Conflicts deeper in the tree were more likely to be correctly inferred by coalescent methods and EPSY. Across all datasets, the mean height of the conflict was greatest for cases where EPSY recovered the true tree (33,648,306 generations), followed coalescent methods (28,000,286 generations) and then concatenation (22,541,833 generations) (Fig.7). Furthermore, the maximum height of a conflicting edge (i.e., closer to the root) was also greatest in cases where EPSY recovers the true tree (99,589,310 generations), followed by coalescent (99,112,112 generations), followed by concatenation (77,920,906 generations). Because investigating conflict height would likely depend on the height of the tree, we analyzed the datasets with shorter root height (100 million generations, datasets A-D) and larger root height (200 million generations, datasets E-H) separately. The mean conflict heights of the shallower datasets showed a similar pattern as when all datasets were included (EPSY 25,155,816 generations; coalescent 23,838,152; concatenation 18,634,964), but the maximum conflict heights were very similar (EPSY 50,000,000 generations; coalescent 50,000,000; concatenation 48,923,774) (Supplemental Fig. S7A). In the deeper datasets, the decreasing EPSY-coalescent-concatenation trend held for the mean conflict height (EPSY 39,798,040 generations; coalescent 30,775,043; concatenation 24,885,955), and there was a discrepancy in the maximum height as when all datasets were included (Supplemental Fig. S7B).

**Figure 7.**
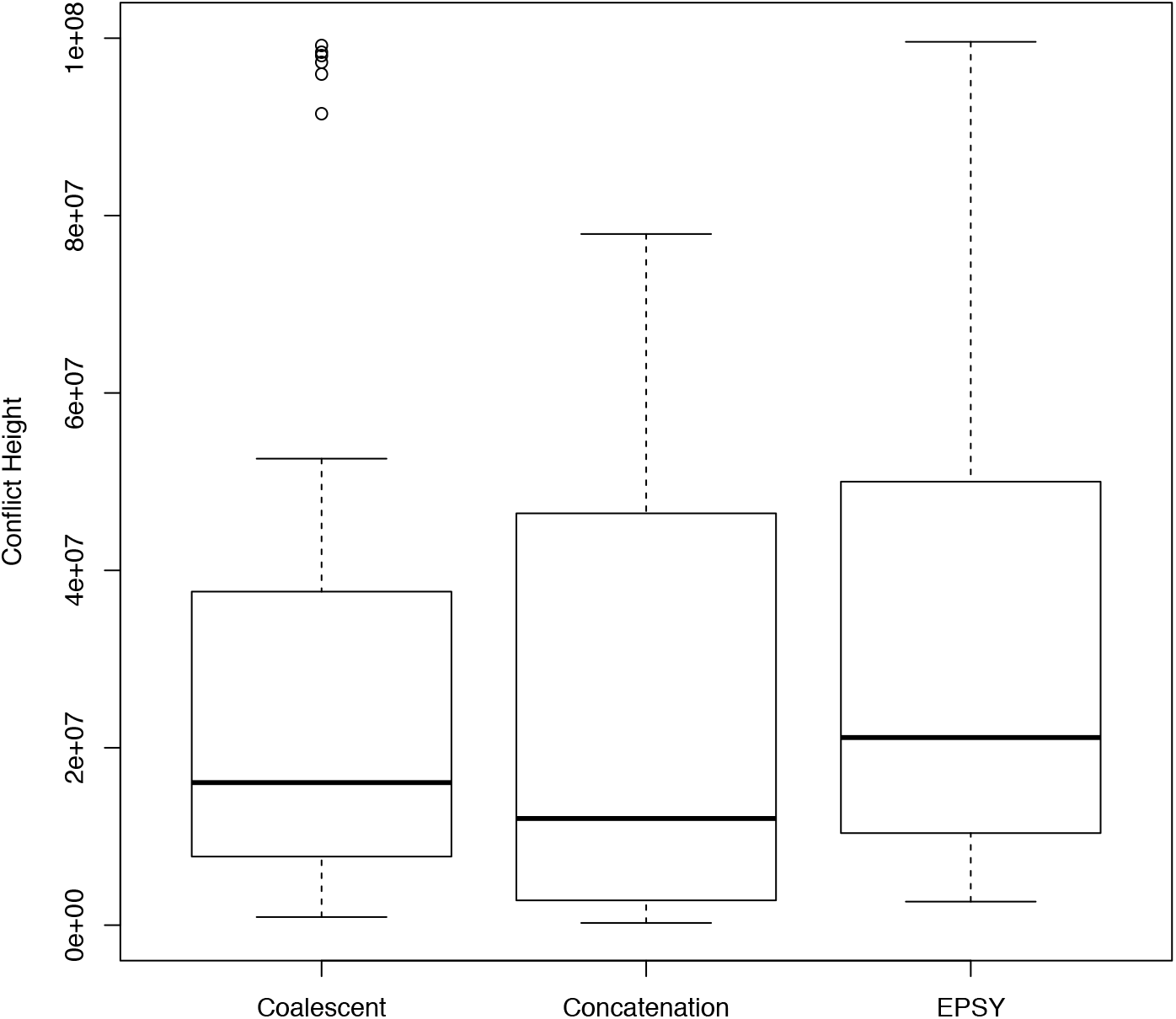
Box-and-whisker plots displaying the height of a conflict for each type of inference method: coalescent, concatenation, and EPSY.

### Empirical application to bird phylogeny

There were numerous conflicts between the coalescent and concatenation topologies inferred for the bird dataset, which was concordant with conflict reported among studies (e.g., major conflicts between Jarvis et al. (2014) and Prum et al. (2015)). Reddy et al.’s (2017) analysis predominantly favored the Jarvis topology, and attributed topological conflict to differences in data type. The topologies inferred in our analyses were largely similar; several groups were shared between our coalescent and concatenation trees (Fig. 2). We used the terminology from Reddy et al. (2017) for consistency, and labeled the groups as ‘magnificent clades’ i, ii, v, vii. Also following Reddy et al. (2017), we labeled certain groups as ‘clades’ for consistency even when they were non-monophyletic. However, there were some key differences on the backbone of the tree: in the coalescent tree, a major clade that includes groups v+vi+vii was sister to a group that includes clades ii and iv, whereas the major clade containing groups v+vi+vii was sister to a clade that includes group i in the concatenation tree. Because our study was interested in cases where ILS is the cause of conflict, we focused on nodes with documented high levels of ILS along the backbone (Suh et al. 2015, Reddy et al. 2017; Fig. 2). We analyzed two conflicting nodes in the backbone of the two competing phylogenies with high levels of ILS in previous studies (Fig. 2). In each of the cases, the results from EPSY, including both the difference in likelihood scores and the number genes, favored the coalescent topology over the concatenation topology by a considerable margin: for both contentious nodes, the sum of differences in likelihood scores was approximately four to seven times larger for genes that supported the coalescent topology, and substantially more genes supported the coalescent topology (Table 2, Supplemental Tables S1 & S2). Removing genes considered non-significant based on the >2 lnL criterion had virtually no effect on the interpretation of the results (Table 2).

**Table 2.** Conflicts along the backbone of the bird phylogeny were compared using EPSY. The number of genes supporting each alternative relationship and difference in log-likelihood (lnL) scores between the two competing alternatives are shown. Additionally, the number of significant genes supporting each alternative relationship and the sum of lnL differences in significant genes is shown. In each of these conflicts, the exclusion of non-significant genes did not qualitatively change the conclusion (see Supplemental Tables S1 & S2 for lnL scores for each gene).

| Conflict | Bipartitions | Alternative | Number genes | Sum lnL difference | Number significant genes | Sum lnL difference: significant genes |
| --- | --- | --- | --- | --- | --- | --- |
| 1 | (i, v, vi, vii, *Charadius*, *Eurypyga*) \| (ii, iv, *Opisthocomus*, *Phaethon*, *Balearica*, Fowl, Flightless) | concatenation | 15 | 128.73741 | 13 | 127.231174 |
|  | (i) \| (ii, iv, v, vi, vii, *Charadius*, *Phaethon*, *Balearica*, *Eurypyga*, *Opisthocomus*, Fowl, Flightless) | coalescent | 44 | 992.35697 | 41 | 989.696992 |
| 2 | (ii, iv, *Opisthocomus*, *Phaethon*, *Balearica*) \| (i, v, vi, vii, *Charadius*, *Eurypyga*, Fowl, Flightless) | concatenation | 21 | 187.03451 | 18 | 183.493725 |
|  | (ii, iv, v, vi, vii, *Charadius*, *Phaethon*, *Balearica*, *Eurypyga*, *Opisthocomus*) \| (i, Fowl, Flightless ) | coalescent | 38 | 779.76696 | 36 | 779.113795 |

We also analyzed the relationship of *Eurypyga* and *Phaethon*, which according to the literature has been resolved through the use of genomic data. The accepted relationship in the recent Neoaves phylogeny was that the orders Eurypygiformes and Phaethontiformes are sister to one another. These two orders were not considered sister to one another well into the genomic era (e.g., in a morphological review (Livezey & Zusi 2007), or in the first pass at a Neoaves phylogeny based on a multilocus dataset (Hackett et al. 2008)). However, subsequently this relationship was inferred repeatedly with different data types and inference methods, frequently with very strong support from Bayesian posterior probabilities and/or bootstrap values (McCormack et al. 2013, Jarvis et al. 2014, Mirarab et al. 2014a, Prum et al. 2015, Suh et al. 2015, Suh 2016, Reddy et al. 2017, Zhang et al. 2018). Notably, neither of our analyses indicated that *Eurypyga helias* was sister to *Phaethon lepturus* (Fig. 2). Therefore, we used EPSY to compare three possible topologies: the two relationships that we inferred (coalescent, concatenation; Fig. 2) and the sister relationship that was overwhelmingly supported in the literature. EPSY was virtually equivocal; the most genes and the sum of log-likelihood score differences supported the concatenation topology (*Phaeton* - *Balearica* sister) inferred in our study (Table 3, Supplemental Table S3). However, when non-significant genes were excluded, most genes favored a *Eurypyga* - *Phaethon* sister relationship, although the sum of log-likelihood score differences still favored the concatenation relationship (Table 3). Regardless, the EPSY results show that each alternative relationship is strongly supported by a substantial number (>10) of genes (Table 3, Supplemental Table S3). This result highlighted how EPSY is valuable for evaluating how confident we should be in existing phylogenetic hypotheses--even those with strong bootstrap and/or posterior probability support.

**Table 3.** Three conflicting alternative relationships tested to investigate whether *Phaethon* + *Eurypyga* is monophyletic. The first topology was inferred by concatenation methods, the second topology was inferred by coalescent methods, and the third topology (*Phaethon* + *Eurypyga* monophyletic) is found in many other studies in the literature. EPSY favored the concatenation topology based on number of genes and likelihood scores but favored the third topology based on number of significant genes, although the concatenation topology was still favored based on the sum of log-likelihood (lnL) scores for significant genes. Examinations of individual genes identify some outlier genes that may be disproportionately affecting the concatenation and coalescent analyses in the present study (see Supplemental Table S3) and could explain why *Phaethon* + *Eurypyga* monophyly was not strongly supported.

| Conflict | Bipartition | Alternative | Number genes | Sum lnL difference | Number significant genes | Sum lnL difference: significant genes |
| --- | --- | --- | --- | --- | --- | --- |
| Phaethon - Eurypyga | (*Phaethon*, *Balearica*) \| (all other taxa) | concatenation | 22 | 145.002025 | 13 | 136.851909 |
|  | (*Eurypyga*, *Opisthocomus*) \| (all other taxa) | coalescent | 16 | 96.773809 | 13 | 94.501542 |
|  | (*Phaethon*, *Eurypyga*) \| (all other taxa) | previous studies | 21 | 104.829764 | 15 | 100.710471 |

## DISCUSSION

We demonstrate that when phylogenomic inference methods disagree, EPSY is an efficient and useful way to increase confidence in contentious relationships. In conditions (both empirical and simulations) where ILS caused conflict, EPSY was consistent with, or outperformed, coalescent methods. Furthermore, EPSY provides researchers with a means of assessing support in relationships other than posterior probabilities or bootstrapping. EPSY allows us to understand when phylogenetic hypotheses are equivocal, even in cases when bootstraps and/or posterior probabilities strongly favor one hypothesis over another. Specifically, EPSY identifies the number of genes that support each competing relationship and the sum of differences in likelihood scores supporting each relationship—providing two quantitative metrics to assess support (or the lack thereof). We also find that EPSY provides a useful form of data interrogation in cases of conflict among major species hypotheses. Not only did EPSY perform well in ILS simulations, but it enabled investigations of conflict in an empirical dataset. EPSY both provided support for contentious relationships along the high-ILS backbone of the Neoaves phylogeny and revealed an underlying conflict within the tree (i.e., the *Eurypyga-Phaethon* relationship) that was masked by strong support scores using conventional metrics. However, in simulations there were still a large number of cases when coalescent and concatenation topologies disagreed but no method identified the correct tree; this occurred in approximately 45% of cases with conflict, indicating EPSY is no panacea. Moreover, there are regions of parameter space--such as when the shortest branch in the species tree and/or the internal branch adjacent to conflict were short--where EPSY and other methods are prone to supporting an incorrect species tree. We also emphasize that EPSY is not a tree inference method--its strength is comparing existing phylogenomic hypotheses, so researchers must have *a priori* phylogenetic relationships to test. Nevertheless, when competing topologies both have strong support using traditional metrics, EPSY can serve as a useful tiebreaker. In this study, we investigated conflict between coalescent and concatenation phylogenies, but these results are also applicable to any situation where two or more competing species tree hypotheses exist.

### The notion of support in phylogenomics: EPSY is an improved metric for assessing phylogenomic support

Historically, the notion of support in phylogenomics has been based on posterior probabilities (Yang & Rannala 1997) or the non-parametric bootstrapping (Felsenstein 1985). Recent efforts to modernize the bootstrap highlight the need to reconsider phylogenetic support for large datasets (Lemoine et al. 2018). However, the fundamental goal of the updated bootstrap remains the same: to facilitate strong statements about phylogenetic relationships. Using these metrics as a gold standard when assessing relationships is clearly problematic, such as in the case of the sister of angiosperms (Xi et al. 2014). One reasonable approach is to isolate specific alternative phylogenetic hypotheses and identify genes with signal supporting alternative relationships (Shen et al. 2017; Brown & Thomson 2017; Walker et al. 2018, Smith et al. 2019). Our results fit into a growing body of literature addressing how we select and filter genes used for phylogenomic inference (Salichos et al. 2014, Chen et al. 2015, Molloy & Warnow 2018). Some studies have concluded that genes should be filtered and clustered based on the underlying process producing conflict (Knowles et al. 2018) or based on phylogenetic signal (Brown & Thompson 2017). Conversely, one may target nodes and select genes that are useful for resolving the node of interest. A logical extension of this point is that genes should be selected for analysis based on having different signals, rather than a shared overall strong signal (Chen et al. 2015; Shen et al. 2017; Smith et al. 2019). When using a targeted approach, the strategy of Chen et al. (2015) to resolve a contentious relationship showed that the selection of genes to help resolve specific phylogenetic questions, rather than genes selected based on phylogenetic signal, ultimately proving useful for determining the branching order of early land birds and jawed vertebrates.

Gene tree heterogeneity implies that phylogenomic inference will benefit from using a diverse set of genes, each of which have value for resolving different parts of the tree (Molloy & Warnow 2018). The myriad evolutionary processes that differentially impact regions of the genome mean that different genes are better at resolving relationships on different time scales, and therefore in different regions of phylogenetic trees (Chen et al. 2015). In many cases, even if datasets have heterogeneous phylogenetic signals, a large enough dataset may enable correct inference of certain problematic nodes. However, it is clear that this is not a tenable way to proceed because large heterogeneous datasets will likely not correctly resolve every relationship in the tree (Smith et al. 2019). Sometimes massive datasets with heterogeneous signals will correctly resolve problematic nodes, but also confidently infer incorrect relationships elsewhere in the tree (Chen et al. 2015). Using all genes without considering their utility in different regions of the tree may lead to strongly supported relationships deep in the tree, but not in shallow regions, or vice versa (Townsend & Leuenberger 2011; Molloy & Warnow 2018). Recent studies noted that species tree estimation should avoid picking genes selected for overall phylogenetic signal, but instead should use a variety of genes, which are useful for addressing different parts of the tree (Townsend & Leuenberger 2011; Chen et al. 2015, Molloy & Warnow 2018). Rather than filtering genes based on support values at problematic nodes (e.g., bootstrapping or posterior probabilities), which creates biases and statistical issues regarding treatment of alternative hypotheses, we now have a method for assessing how well each gene supports a given relationship, and how much signal a gene has when addressing specific relationships and/or regions of the tree.

### An empirical example of hidden conflict

We used EPSY to compare the relationships reconstructed from concatenation and coalescent analyses involving the *Phaethon-Eurypyga* sister relationship (Fig. 2, Table 3). The results show that EPSY finds higher support for a *Phaethon-Balearica* sister relationship inferred by concatenation based on the number of genes (22 out of 59; 37.3%) and summed difference in log-likelihood scores. However, nearly as many genes supported the *Phaethon-Eurypyga* sister (21 out of 59; 35.6%) and 16 genes (27.1%) support the third relationship inferred by the coalescent (Table 3). If non-significant genes are excluded, the *Phaethon-Eurypyga* sister relationship is narrowly supported based on gene number (15 genes vs. 13 each for concatenation and coalescent; Table 3), but the concatenation relationship is still favored by the summed difference of log-likelihood scores. This example highlights how relationships that are virtually equivocal due to large amounts of underlying conflict can have high support--in many cases with 100% bootstrap support and/or posterior probabilities of 1.0. There is large variation in the ΔlnLscores across the 59 genes, suggesting that certain genes disproportionately favor certain relationships (see Supplemental Information, Supplemental Table 3). The *Phaethon-Eurypyga* relationship is not in the area of the Neoaves phylogeny identified to have high ILS, so we do not know the cause of the conflict. However, using EPSY focused on this node clearly demonstrated that even when relationships have the strongest possible support with existing metrics (i.e., bootstraps), there can be hidden undetected conflict. This empirical example highlights the need for alternative metrics of phylogenomic support.

### Which phylogenomic methods should we use?

Our work highlights nuances of frequently-utilized phylogenomic methods. One massive obstacle to species-tree estimation is long branch attraction, which Roch et al. (2019) note may confound ML analyses when there are finite sequences lengths for the genes used in a phylogenomic analysis. In empirical datasets, clearly the number of sites in a given gene must be finite. A recent simulation study found that many methods are statistically inconsistent (i.e., do not increase the probability of returning a true species tree as the number of loci and sites increases (Bayzid et al. 2015; Roch et al. 2019)). Roch et al. (2019) discovered that when the number of sites per gene is finite, fully partitioned ML is inconsistent and/or positively misleading, and that summary methods can be positively misleading even with no gene tree heterogeneity. Even when all genes evolve under the same model tree, none are statistically consistent (and are sometimes positively misleading) when the number of sites is finite (Roch et al. 2019). Although we can acquire orders of magnitudes more data than we could a quarter century ago, data will always be finite. The adage of the genomic era – as we add more data, surely species tree accuracy must increase – may not hold true. This represents a damning statement for some modern phylogenomic methods, which may only be accurate (i.e., statistically consistent) under unrealistic, impossible conditions (i.e., infinite length of gene sequences).

The behavior of EPSY methods have not been extensively tested; here we demonstrate some of the properties of EPSY, and document the conditions when this method recovers the true tree in comparison to coalescent and concatenation methods. EPSY methods are data centric and make no assumptions about the process or processes that generated the heterogeneity. Here, we demonstrated via simulations that when the source of phylogenetic conflict is known (i.e., ILS) and accounted for in analyses (by using methods that account for ILS, i.e., analyses statistically consistent with the coalescent; ASTRAL), EPSY typically supports the topologies inferred by ASTRAL. Because the simulations modeled high-ILS conditions, and EPSY was qualitatively and quantitatively consistent with the coalescent analyses, we provided evidence that EPSY recovers the true tree when the process generating conflict is known. We also demonstrated that there are certain regions of tree space where no method investigated can recover a true topology; the description of these regions of tree space will be valuable for future researchers. In general, as the length of the shortest branch in a species tree decreased, the length of the internal edge adjacent to the conflict decreased, and the height of the conflict increased, it was more likely that no method would recover the true tree. Furthermore, we demonstrated that in some cases with simulated data, EPSY can support the species tree topology even when both coalescent and concatenation analyses fail to correctly infer it. Investigations with empirical data reinforced findings from the simulations; specifically when ILS is the source of conflict, EPSY consistently favors the tree inference method that accounts for ILS (i.e., coalescence) over one that does not (i.e., concatenation). We emphasize that EPSY is not a tree-inference method and require prior phylogenetic hypotheses. In cases where there are two or more competing hypotheses each supported by data, EPSY is a powerful tool for distinguishing the most likely scenario.

### Portions of the tree where EPSY methods are most valuable

The height of the internal edge adjacent to the conflict of a species tree predicted when certain methods would reliably find the true tree. Importantly, EPSY performed better deeper in phylogenetic trees, which has often been a problem area (e.g., it is frequently difficult to obtain high bootstrap support deep in a phylogeny; Lemoine et al. 2018). Therefore, EPSY is a valuable tool for assessing support for contentious edges deeper in trees. Lanier & Knowles (2015) observed that while coalescent methods rose to popularity because of their ability to perform well for shallow divergences, they also are useful for deeper divergences, although they are computationally intractable as dataset size increased. Concatenation methods are computationally more manageable and generally perform well, although they assume all gene trees share an underlying topology, a feature which causes concatenation to perform poorly when there is pervasive ILS (Kubatko & Degnan 2007). Moreover, both coalescent and concatenation methods are statistically inconsistent and/or positively misleading under conditions common to many empirical datasets (Roch et al. 2019). Here, we document a method with a low computational burden that outperforms both coalescent and concatenation approaches in many regions of simulated phylogenies.

We demonstrated that some properties of a tree, such as the length of the shortest branch in a species tree, or the length of branches adjacent to a conflict, can influence which methods are more likely to accurately infer a contentious relationship. The length of the internal edge adjacent to a conflict was a good predictor of which methods would recover the true species tree. In general, EPSY and coalescent analyses could recover the true topology even as adjacent branches got very long (Fig. 6). In some respects, these results recapitulate the findings of Liu et al. (2015), which reported that long internal branches often confound concatenation methods, but not coalescent methods, in the presence of ILS. We also demonstrate that long internal branches do not confound EPSY results. The height of a coalescent-concatenation conflict also predicted which methods would perform better. These results conform with Lanier & Knowles (2015), which found that coalescent methods are useful for addressing deep divergences.

### High-ILS nodes in the Neoaves phylogeny

Several contentious nodes along the backbone of the Neoaves phylogeny, where there is documented high ILS, have been contentious despite frequent interrogation. Here, we demonstrated a means of providing support for contentious relationships that is not dependent on bootstrap scores or posterior probabilities, both of which can be positively misleading (e.g., Hillis & Bull 1993, Suzuki et al. 2002). EPSY showed strong support for the coalescent topology in both cases in our empirical example. We emphasize that this is an example of how EPSY can be used in a typical dataset where the goal is to resolve several specific relationships, but we do not make strong claims about the phylogenetic placement of taxa in the bird phylogeny. As noted in many previous studies, there can be many reasons (e.g., data type; Reddy et al. 2017) that one would want to carefully curate a dataset in different ways. Nevertheless, assuming our treatment of the 59-gene, 48-taxon dataset followed best practices that a researcher would use to implement a coalescent- or concatenation-based tree reconstruction, we show clear support for the coalescent topology at each of the contentious edges. We emphasize that we selected this dataset because the backbone is notoriously difficult to resolve, so much so that some treat it as a hard polytomy (Suh 2016), and because of the high degree of ILS documented along the backbone. We are not making strong statements about these specific relationships, but we have provided strong evidence that EPSY is consistent with the coalescent when the cause of conflict (i.e., ILS) is known and accounted for in analyses, and have documented the underlying uncertainty at key nodes in the tree.

### Recommendations to researchers

Contentious nodes may often be resolved with conflicting resolutions using alternative methods, each with high support leaving researchers uncertain how to proceed with confidence (Pease et al. 2018). Typical solutions include increased taxon sampling (e.g., Zwickl & Hillis 2002), adding genetic data, or identifying potential biases that impact phylogenetic inference (e.g., Reddy et al. 2017). However, for some key charismatic clades, data has increased rapidly yet these conflicts remain. Some relationships, such as the branching order of ancestors of jawed vertebrates, birds, and metazoa will remain contentious using established phylogenomic methods (Near 2009, Janvier 2010, Arcila et al. 2017, Miyashita et al. 2019). There are several instances where recovered conflict was the result of systematic error and should be removed (Jeffroy et al. 2006, Philippe et al. 2011). When conflict is the result of biological processes and not systematic error, however, we should not expect the addition of data to abate conflict. We need to consider whether some contentious relationships can be resolved, and whether it is worth the monumental effort to collect more data with the goal of resolving them when other regions of the Tree of Life remain unexplored. In fact, if the source of conflict is biological, we would expect the increase in data to result in the recovery of more conflict. We found that with simulated high-ILS datasets, there are portions of phylogenetic tree parameter space where one or both methods fail to recover the true tree. We stress that we are not making broad generalizations about the utility of the coalescent and concatenation methods used in this study, but rather note that under the set of parameters investigated, EPSY outperforms both methods when the goal is investigating a specific relationship. Many phylogenetic relationships in the Tree of Life may have virtually equivocal resolutions, and it may be preferable to acknowledge this uncertainty, as opposed to searching for ways to filter data to make opposing, but strongly supported statements. We suggest that the conflict itself represents the underlying evolutionary history of those genes within those lineages. For these highly contentious and conflicting areas of the Tree of Life, we suggest that EPSY should be used in concert with other inference methods to investigate nodes with uncertain resolution and/or large amounts of conflict. Even if these analyses do not resolve the relationship, the underlying data will be better explored and researchers may feel more confident in the lack of ability to resolve the particular relationship.

### Conclusions

Biological processes shape the Tree of Life. These events (e.g., rapid radiations, population contractions and expansions, adaptation, mass extinctions, and genome evolution) have left complex and conflicting signals that underlie the genomic data we use for phylogenomic inference. Increasingly larger datasets have been gathered to help better understand conflict in hopes of resolving difficult areas across the Tree of Life. Here, we show that, despite these large datasets, no current phylogenomic inference method is up to the task of solving *all* difficult relationships even when the process causing conflict is known. However, our analyses of how coalescent, concatenation, and EPSY methods behave in the presence of conflict yielded several key insights that should guide future research. When competing topologies inferred using different genomic datasets or different inference methods both have strong support using traditional metrics, EPSY serves as a useful tiebreaker. Furthermore, when the objective is to understand why a relationship of interest is contentious, EPSY can quantify the amount of underlying conflict and provide the researcher with a nuanced view of the gene tree conflict associated with the relationship of interest. We find value in all current phylogenetic methods and encourage researchers pursuing questions involving large dataset to continue interrogating the data using multiple analyses. We encourage a more agnostic approach to phylogenomic conflict, where the goal is to understand and quantify the underlying conflict, as opposed to manipulating data to optimize our ability to make strong opposing statements. Such an approach would force us to reconsider how confident we should be in relationships that were previously considered unequivocal, and will better guide future analyses investigating the biological causes of phylogenomic conflict. Finally, we contend that further examination of instances where no method is capable of resolving relationships is warranted.

## Acknowledgements

We thank Ning Wang for many valuable comments that helped improve the manuscript.

