## Supplemental Table 2 for "When species trees disagree: an approach consistent with the coalescent that quantifies phylogenomic support for contentious relationships"

**Table S2.** Log-likelihood (lnL) scores for each gene addressing the alternative relationships from conflict 2 in Table 2. Genes are sorted based on which alternative they support (concatenation vs. coalescent) and then by magnitude of difference in lnL score.

| **Gene** | **lnL score-concatenation** | **lnL score-coalescent** | **Best one** | **Difference lnL scores** |
| --- | --- | --- | --- | --- |
| aldob.i6.phy | -10798.17723 | -10826.45893 | Concatenation | 28.281693 |
| fgb4.phy | -11839.44686 | -11866.35438 | Concatenation | 26.907515 |
| ngf.phy | -7443.136374 | -7460.572865 | Concatenation | 17.436491 |
| ppp2cb.phy | -14303.46058 | -14320.31782 | Concatenation | 16.85724 |
| fgb67.i6.phy | -3246.035956 | -3259.808176 | Concatenation | 13.77222 |
| fgb5.phy | -10397.70801 | -10410.44389 | Concatenation | 12.735878 |
| eef2.e.phy | -3561.178977 | -3572.953609 | Concatenation | 11.774632 |
| eef2.i5.phy | -10069.61752 | -10079.53607 | Concatenation | 9.918555 |
| park7.phy | -14670.6889 | -14680.51242 | Concatenation | 9.823521 |
| eef2.i6.phy | -6191.855596 | -6198.542605 | Concatenation | 6.687009 |
| per2.phy | -11373.30541 | -11378.29098 | Concatenation | 4.985573 |
| irf2.phy | -8648.315521 | -8653.019176 | Concatenation | 4.703655 |
| cltc.e.phy | -2053.338081 | -2057.432424 | Concatenation | 4.094343 |
| vdac2.phy | -8826.887619 | -8830.753669 | Concatenation | 3.86605 |
| gars.phy | -11910.61857 | -11914.30512 | Concatenation | 3.686544 |
| hoxa3.i1.phy | -25431.69415 | -25434.64862 | Concatenation | 2.954468 |
| pde6b.phy | -12882.9523 | -12885.57752 | Concatenation | 2.625223 |
| aldob.e.phy | -4255.252246 | -4257.635361 | Concatenation | 2.383115 |
| hnrnpa2b1.phy | -7569.715509 | -7571.213721 | Concatenation | 1.498212 |
| fgb67.e.phy | -3194.697311 | -3195.72151 | Concatenation | 1.024199 |
| myc.e.phy | -3654.750229 | -3655.768605 | Concatenation | 1.018376 |
| pcbd1.i3.phy | -11026.82197 | -10927.46331 | Coalescent | 99.358654 |
| cltc.i6.phy | -16087.32322 | -15988.08954 | Coalescent | 99.233678 |
| cltc.i7.phy | -13289.83498 | -13237.07045 | Coalescent | 52.764525 |
| fgb67.i7.phy | -16923.62188 | -16872.12279 | Coalescent | 51.499092 |
| pcbd1.i2.phy | -9467.558081 | -9423.615034 | Coalescent | 43.943047 |
| naa60.phy | -17210.60694 | -17167.60238 | Coalescent | 43.00456 |
| clock6.phy | -12374.06378 | -12331.57053 | Coalescent | 42.493248 |
| csnk1e.phy | -10954.95655 | -10918.71812 | Coalescent | 36.23843 |
| eef2.i8.phy | -12671.87773 | -12648.497 | Coalescent | 23.380727 |
| bdnf.phy | -4008.497619 | -3987.581473 | Coalescent | 20.916146 |
| rho.i1.phy | -18782.73861 | -18761.99595 | Coalescent | 20.742662 |
| irf1.phy | -22245.65288 | -22225.14742 | Coalescent | 20.505459 |
| musk.i3.phy | -9950.47389 | -9930.974539 | Coalescent | 19.499351 |
| hmgn45.i4.phy | -10357.10082 | -10338.58098 | Coalescent | 18.519839 |
| myc.i2.phy | -6055.165451 | -6039.089296 | Coalescent | 16.076155 |
| cltcl1.phy | -8522.126914 | -8506.45427 | Coalescent | 15.672644 |
| paxip1.phy | -8754.644913 | -8739.972172 | Coalescent | 14.672741 |
| ntf3.phy | -5534.729047 | -5520.449996 | Coalescent | 14.279051 |
| aldob.i5.phy | -5441.108898 | -5427.121105 | Coalescent | 13.987793 |
| eif5.phy | -12682.43789 | -12669.60621 | Coalescent | 12.831676 |
| tgfb2.phy | -10173.20708 | -10161.03522 | Coalescent | 12.171862 |
| psma2.phy | -8397.675071 | -8385.862316 | Coalescent | 11.812755 |
| egr1u.phy | -4434.649127 | -4424.410651 | Coalescent | 10.238476 |
| eef2.i7.phy | -5044.128847 | -5035.813969 | Coalescent | 8.314878 |
| cryaa.e.phy | -1398.606983 | -1390.399997 | Coalescent | 8.206986 |
| hmgn45.i3.phy | -9575.658371 | -9568.51509 | Coalescent | 7.143281 |
| aco1.phy | -20181.6511 | -20174.96348 | Coalescent | 6.687623 |
| cyp19a1.5u.phy | -5562.747168 | -5556.787724 | Coalescent | 5.959444 |
| egr1c.phy | -10555.89423 | -10550.3875 | Coalescent | 5.506732 |
| aldob.i4.phy | -2871.356383 | -2865.90666 | Coalescent | 5.449723 |
| vim.phy | -11888.89068 | -11885.03771 | Coalescent | 3.852962 |
| sept2.phy | -8856.535479 | -8853.232887 | Coalescent | 3.302592 |
| clocku.phy | -5588.854194 | -5585.784322 | Coalescent | 3.069872 |
| srsf3.phy | -8338.767945 | -8335.844751 | Coalescent | 2.923194 |
| myc.3u.phy | -2899.770278 | -2897.130371 | Coalescent | 2.639907 |
| calb1.phy | -9285.940869 | -9283.726839 | Coalescent | 2.21403 |
| cryaa.i1.phy | -20153.53025 | -20153.03089 | Coalescent | 0.499355 |
| rho.e.phy | -4069.957855 | -4069.804046 | Coalescent | 0.153809 |
