## Supplemental Table 1 for "When species trees disagree: an approach consistent with the coalescent that quantifies phylogenomic support for contentious relationships"

**Table S1.** Log-likelihood (lnL) scores for each gene addressing the alternative relationships from conflict 1 in Table 2. Genes are sorted based on which alternative they support (concatenation vs. coalescent) and then by magnitude of difference in lnL score.

| **Gene** | **lnL score: concatenation** | **lnL score: coalescent** | **Best one** | **Difference lnL scores** |
| --- | --- | --- | --- | --- |
| aldob.i6.phy | -10794.77836 | -10828.27891 | Concatenation | 33.500551 |
| ngf.phy | -7445.584793 | -7460.881412 | Concatenation | 15.296619 |
| eef2.i5.phy | -10071.85983 | -10086.0803 | Concatenation | 14.220475 |
| fgb67.e.phy | -3186.364921 | -3200.343325 | Concatenation | 13.978404 |
| vdac2.phy | -8820.794119 | -8831.80688 | Concatenation | 11.012761 |
| eef2.e.phy | -3565.332708 | -3575.964529 | Concatenation | 10.631821 |
| park7.phy | -14678.22454 | -14684.81886 | Concatenation | 6.594323 |
| per2.phy | -11371.73128 | -11377.25995 | Concatenation | 5.528671 |
| gars.phy | -11908.88981 | -11913.66782 | Concatenation | 4.778003 |
| aldob.e.phy | -4251.146101 | -4254.329047 | Concatenation | 3.182946 |
| fgb67.i6.phy | -3244.381872 | -3247.408327 | Concatenation | 3.026455 |
| fgb5.phy | -10397.7136 | -10400.64949 | Concatenation | 2.935891 |
| hnrnpa2b1.phy | -7570.224365 | -7572.768619 | Concatenation | 2.544254 |
| cltc.e.phy | -2058.669636 | -2059.756186 | Concatenation | 1.08655 |
| eef2.i6.phy | -6194.237653 | -6194.65734 | Concatenation | 0.419687 |
| cltc.i6.phy | -16085.1329 | -15982.02572 | Coalescent | 103.107179 |
| pcbd1.i3.phy | -11018.66747 | -10922.05834 | Coalescent | 96.609133 |
| fgb67.i7.phy | -16923.26865 | -16857.29703 | Coalescent | 65.971621 |
| cltc.i7.phy | -13288.30266 | -13222.88287 | Coalescent | 65.419788 |
| naa60.phy | -17217.30921 | -17165.44565 | Coalescent | 51.863555 |
| clock6.phy | -12371.58538 | -12331.75904 | Coalescent | 39.826347 |
| pcbd1.i2.phy | -9463.603896 | -9425.90183 | Coalescent | 37.702066 |
| ppp2cb.phy | -14328.11514 | -14295.41167 | Coalescent | 32.703462 |
| hoxa3.i1.phy | -25435.63144 | -25403.05145 | Coalescent | 32.579995 |
| paxip1.phy | -8763.602348 | -8736.965682 | Coalescent | 26.636666 |
| sept2.phy | -8866.442467 | -8840.432504 | Coalescent | 26.009963 |
| rho.i1.phy | -18783.60414 | -18758.77038 | Coalescent | 24.833757 |
| musk.i3.phy | -9949.178356 | -9924.376865 | Coalescent | 24.801491 |
| psma2.phy | -8402.478658 | -8378.027096 | Coalescent | 24.451562 |
| csnk1e.phy | -10948.62303 | -10927.44901 | Coalescent | 21.174022 |
| aldob.i5.phy | -5442.848159 | -5421.727759 | Coalescent | 21.1204 |
| srsf3.phy | -8339.251694 | -8320.600199 | Coalescent | 18.651495 |
| eef2.i8.phy | -12662.70235 | -12644.38268 | Coalescent | 18.319667 |
| tgfb2.phy | -10172.51268 | -10154.89108 | Coalescent | 17.621595 |
| irf1.phy | -22242.60085 | -22225.38051 | Coalescent | 17.220341 |
| irf2.phy | -8658.016936 | -8640.815348 | Coalescent | 17.201588 |
| ntf3.phy | -5533.565177 | -5517.389218 | Coalescent | 16.175959 |
| hmgn45.i3.phy | -9584.191115 | -9568.141218 | Coalescent | 16.049897 |
| hmgn45.i4.phy | -10354.19912 | -10338.40635 | Coalescent | 15.792772 |
| bdnf.phy | -4001.987037 | -3986.383132 | Coalescent | 15.603905 |
| cryaa.e.phy | -1403.791608 | -1390.333417 | Coalescent | 13.458191 |
| myc.i2.phy | -6054.177853 | -6040.803935 | Coalescent | 13.373918 |
| cryaa.i1.phy | -20164.99526 | -20152.55624 | Coalescent | 12.439019 |
| eef2.i7.phy | -5046.006322 | -5034.410927 | Coalescent | 11.595395 |
| egr1u.phy | -4434.772825 | -4423.217298 | Coalescent | 11.555527 |
| cltcl1.phy | -8523.999121 | -8512.681419 | Coalescent | 11.317702 |
| myc.e.phy | -3662.935053 | -3651.62183 | Coalescent | 11.313223 |
| pde6b.phy | -12882.3081 | -12872.76143 | Coalescent | 9.546673 |
| cyp19a1.5u.phy | -5565.141591 | -5556.578168 | Coalescent | 8.563423 |
| clocku.phy | -5591.467454 | -5583.855213 | Coalescent | 7.612241 |
| rho.e.phy | -4068.955758 | -4062.202953 | Coalescent | 6.752805 |
| egr1c.phy | -10554.7443 | -10548.58162 | Coalescent | 6.162679 |
| aldob.i4.phy | -2871.459923 | -2866.424336 | Coalescent | 5.035587 |
| myc.3u.phy | -2899.403342 | -2894.487908 | Coalescent | 4.915434 |
| vim.phy | -11881.45265 | -11876.59658 | Coalescent | 4.856077 |
| calb1.phy | -9282.884659 | -9279.133787 | Coalescent | 3.750872 |
| eif5.phy | -12666.51543 | -12664.99162 | Coalescent | 1.523813 |
| aco1.phy | -20171.02316 | -20169.99555 | Coalescent | 1.027608 |
| fgb4.phy | -11838.98601 | -11838.87745 | Coalescent | 0.108557 |
