## Supplemental Table 3 for "When species trees disagree: an approach consistent with the coalescent that quantifies phylogenomic support for contentious relationships"

**Table S3.** Log-likelihood (lnL) scores for each gene addressing the three alternative relationships from Table 3. Genes are sorted based on which alternative they support (concatenation vs. coalescent vs. previous hypothesis) and then by magnitude of difference in lnL score.

| **Gene** | **lnL score-concatenation** | **lnL score-coalescent** | **lnL score-previous** | **Best one** | **Difference lnL scores** |
| --- | --- | --- | --- | --- | --- |
| fgb5.phy | -10398.96201 | -10439.57375 | -10436.38261 | Concatenation | 37.420606 |
| fgb4.phy | -11835.67399 | -11862.83303 | -11860.72468 | Concatenation | 25.050686 |
| eef2.i6.phy | -6161.544283 | -6182.668187 | -6172.891667 | Concatenation | 11.347384 |
| aldob.e.phy | -4230.891187 | -4242.327799 | -4241.461404 | Concatenation | 10.570217 |
| cltc.i7.phy | -13208.52305 | -13218.36772 | -13255.54964 | Concatenation | 9.844673 |
| egr1u.phy | -4421.993968 | -4430.615454 | -4434.517734 | Concatenation | 8.621486 |
| cryaa.i1.phy | -20125.3813 | -20133.81821 | -20161.96303 | Concatenation | 8.436908 |
| park7.phy | -14652.54502 | -14671.75864 | -14658.65987 | Concatenation | 6.114856 |
| srsf3.phy | -8320.594077 | -8324.863768 | -8332.779227 | Concatenation | 4.269691 |
| hoxa3.i1.phy | -25365.21279 | -25380.32215 | -25369.439 | Concatenation | 4.226203 |
| eef2.e.phy | -3527.433992 | -3531.429696 | -3540.572066 | Concatenation | 3.995704 |
| hmgn45.i3.phy | -9566.495957 | -9570.958521 | -9570.155761 | Concatenation | 3.659804 |
| eef2.i7.phy | -5002.98445 | -5006.278141 | -5009.15957 | Concatenation | 3.293691 |
| naa60.phy | -17165.76029 | -17169.44927 | -17167.75657 | Concatenation | 1.996282 |
| fgb67.e.phy | -3161.830388 | -3163.708776 | -3163.872418 | Concatenation | 1.878388 |
| bdnf.phy | -3976.116402 | -3977.539725 | -3979.51157 | Concatenation | 1.423323 |
| rho.e.phy | -4023.582381 | -4024.349521 | -4039.2616 | Concatenation | 0.76714 |
| psma2.phy | -8361.145333 | -8363.436212 | -8361.766054 | Concatenation | 0.620721 |
| musk.i3.phy | -9918.45466 | -9918.917578 | -9924.772559 | Concatenation | 0.462918 |
| rho.i1.phy | -18758.49948 | -18758.94683 | -18768.449 | Concatenation | 0.447354 |
| pde6b.phy | -12851.40417 | -12854.81565 | -12851.81508 | Concatenation | 0.410908 |
| cyp19a1.5u.phy | -5554.232273 | -5554.375355 | -5555.671393 | Concatenation | 0.143082 |
| irf2.phy | -8636.753272 | -8623.133803 | -8636.522759 | Coalescent | 13.388956 |
| vim.phy | -11869.65152 | -11856.81887 | -11877.52107 | Coalescent | 12.832645 |
| clocku.phy | -5566.535702 | -5554.400838 | -5565.260023 | Coalescent | 10.859185 |
| fgb67.i6.phy | -3257.724613 | -3240.355136 | -3249.369428 | Coalescent | 9.014292 |
| csnk1e.phy | -10931.68909 | -10913.03073 | -10921.61993 | Coalescent | 8.589195 |
| per2.phy | -11339.84792 | -11332.19053 | -11346.39221 | Coalescent | 7.657382 |
| cryaa.e.phy | -1378.215269 | -1365.730367 | -1371.945049 | Coalescent | 6.214682 |
| ngf.phy | -7423.844388 | -7417.860984 | -7424.133539 | Coalescent | 5.983404 |
| hmgn45.i4.phy | -10341.71253 | -10320.57151 | -10325.64953 | Coalescent | 5.078018 |
| sept2.phy | -8846.816383 | -8837.804674 | -8842.461404 | Coalescent | 4.65673 |
| myc.i2.phy | -6042.862221 | -6037.331701 | -6041.809262 | Coalescent | 4.477561 |
| paxip1.phy | -8743.863945 | -8739.403078 | -8742.915108 | Coalescent | 3.51203 |
| egr1c.phy | -10529.90499 | -10527.66753 | -10532.68589 | Coalescent | 2.237462 |
| cltc.i6.phy | -15963.90138 | -15962.8131 | -16018.5908 | Coalescent | 1.088281 |
| aldob.i4.phy | -2847.5501 | -2842.111377 | -2842.792845 | Coalescent | 0.681468 |
| cltc.e.phy | -2038.470757 | -2037.801329 | -2038.303847 | Coalescent | 0.502518 |
| pcbd1.i3.phy | -10926.11504 | -10925.35677 | -10910.16951 | Previous Hypothesis | 15.187257 |
| cltcl1.phy | -8498.52347 | -8491.643082 | -8477.435708 | Previous Hypothesis | 14.207374 |
| hnrnpa2b1.phy | -7543.38599 | -7545.371902 | -7532.336652 | Previous Hypothesis | 11.049338 |
| eef2.i5.phy | -10061.44966 | -10060.19543 | -10049.58389 | Previous Hypothesis | 10.611541 |
| fgb67.i7.phy | -16945.14344 | -16869.40091 | -16859.86975 | Previous Hypothesis | 9.531162 |
| pcbd1.i2.phy | -9409.768138 | -9410.609782 | -9402.446948 | Previous Hypothesis | 7.32119 |
| myc.3u.phy | -2870.69652 | -2863.407742 | -2856.956483 | Previous Hypothesis | 6.451259 |
| aldob.i5.phy | -5414.940159 | -5418.236523 | -5409.663341 | Previous Hypothesis | 5.276818 |
| aldob.i6.phy | -10784.10167 | -10789.02928 | -10780.26262 | Previous Hypothesis | 3.839057 |
| tgfb2.phy | -10136.85164 | -10136.66351 | -10133.00726 | Previous Hypothesis | 3.656254 |
| clock6.phy | -12318.68119 | -12339.9796 | -12315.74886 | Previous Hypothesis | 2.932329 |
| eef2.i8.phy | -12602.21931 | -12598.04622 | -12595.1143 | Previous Hypothesis | 2.931918 |
| aco1.phy | -20133.9839 | -20156.83979 | -20131.12793 | Previous Hypothesis | 2.855965 |
| myc.e.phy | -3643.387383 | -3653.636565 | -3640.565114 | Previous Hypothesis | 2.822269 |
| vdac2.phy | -8795.360967 | -8793.132702 | -8791.095962 | Previous Hypothesis | 2.03674 |
| irf1.phy | -22204.6321 | -22230.19981 | -22202.72732 | Previous Hypothesis | 1.904788 |
| gars.phy | -11885.31534 | -11884.66469 | -11883.99815 | Previous Hypothesis | 0.666547 |
| calb1.phy | -9263.757126 | -9270.469485 | -9263.186517 | Previous Hypothesis | 0.570609 |
| ntf3.phy | -5515.129959 | -5506.272876 | -5505.837144 | Previous Hypothesis | 0.435732 |
| ppp2cb.phy | -14274.09238 | -14282.3232 | -14273.71697 | Previous Hypothesis | 0.375408 |
| eif5.phy | -12673.84505 | -12636.23417 | -12636.06796 | Previous Hypothesis | 0.166209 |
