## Supplement Legends for "When species trees disagree: an approach consistent with the coalescent that quantifies phylogenomic support for contentious relationships"

\## Supplemental Tables

Table S1. Log-likelihood (lnL) scores for each gene addressing the alternative relationships from conflict 1 in Table 2. Genes are sorted based on which alternative they support (concatenation vs. coalescent) and then by magnitude of difference in lnL score.

Table S2. Log-likelihood (lnL) scores for each gene addressing the alternative relationships from conflict 2 in Table 2. Genes are sorted based on which alternative they support (concatenation vs. coalescent) and then by magnitude of difference in lnL score.

Table S3. Log-likelihood (lnL) scores for each gene addressing the three alternative relationships from Table 3. Genes are sorted based on which alternative they support (concatenation vs. coalescent vs. previous hypothesis) and then by magnitude of difference in lnL score.

\## Supplemental Figures

Figure S1. Example simulated species trees for each of the eight datasets.  See Table 1 for the parameterization of each of the datasets.

Figure S2. The number of species trees, out of 50 total, where there is conflict between the coalescent and concatenation topologies for each dataset.

Figure S3. Box-and-whisker plots describing the normalized average Robinson-Foulds distances between each species tree and the true gene trees in the species tree are displayed for all eight datasets.

Figure S4. For all eight datasets, the proportion of species trees where no method can correctly identify the true topology.

Figure S5. A) For the datasets with very high ILS (C, D, G, H; based on R-F distances), box-and-whisker plots display the proportion of conflicts that are correctly inferred (left) by each method, and the proportion never correctly inferred (right). B) For the datasets with high ILS (A, B, E, F), box-and-whisker plots display the proportion of conflicts that are correctly inferred (left) by each method, and the proportion never correctly inferred (right).

Figure S6. The mean (+/- standard error) length of the shortest branch in the species tree across all datasets for cases where there is conflict, no conflict, or conflict so extreme that no method infers the correct phylogeny.

Figure S7. A) For the four datasets with shorter root height, box-and-whisker plots show the range of conflict heights for each inference method. B) For the datasets with larger root height, the range of conflict heights for each inference method is displayed.
