## Supplementary figures and images for "When species trees disagree: an approach consistent with the coalescent that quantifies phylogenomic support for contentious relationships"

### Supplemental Figure 1

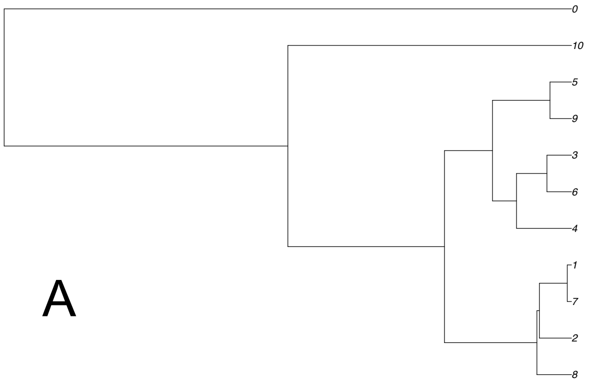

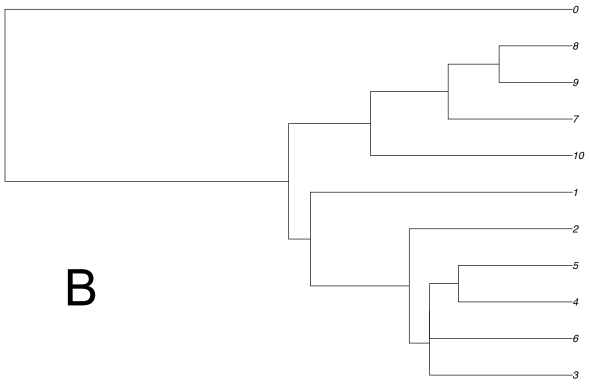

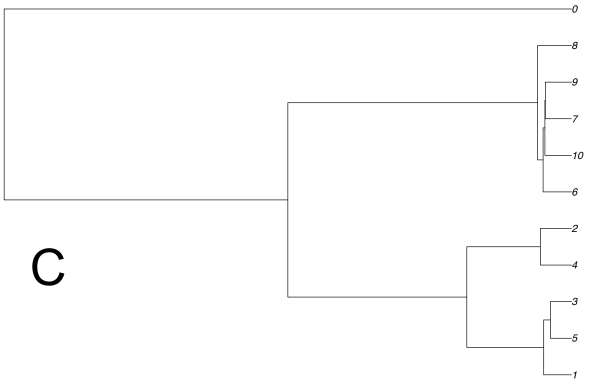

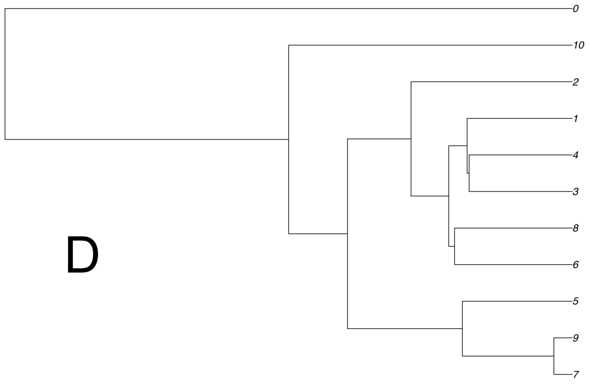

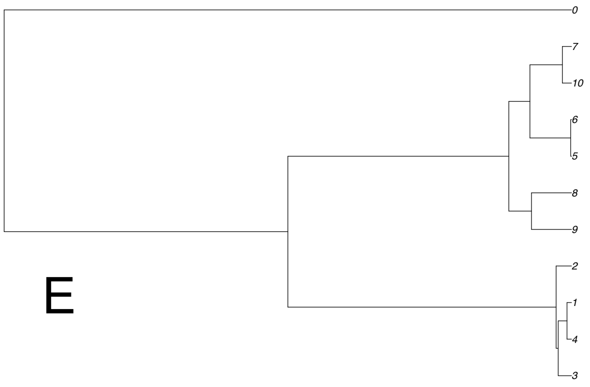

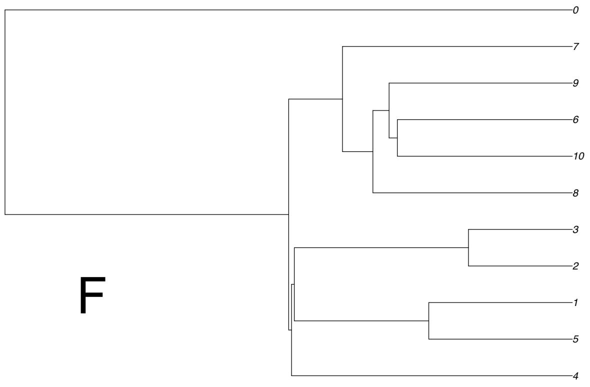

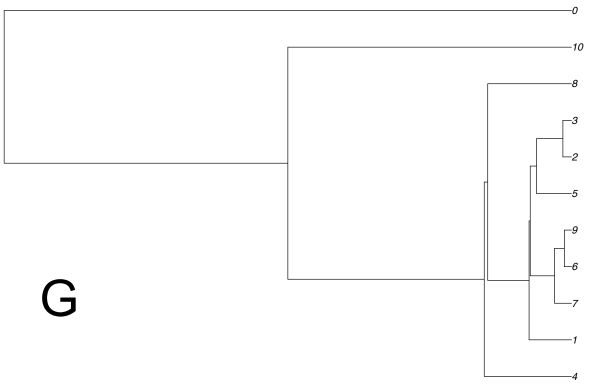

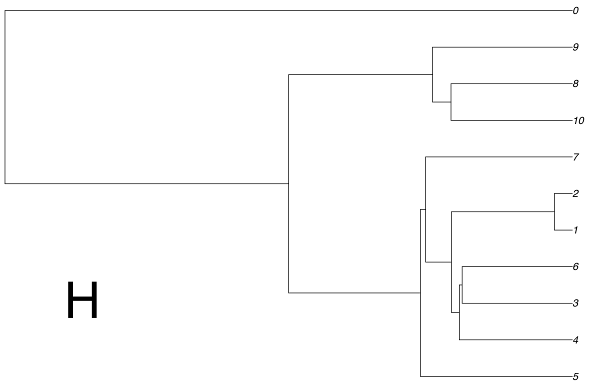

### Supplemental Figure 2

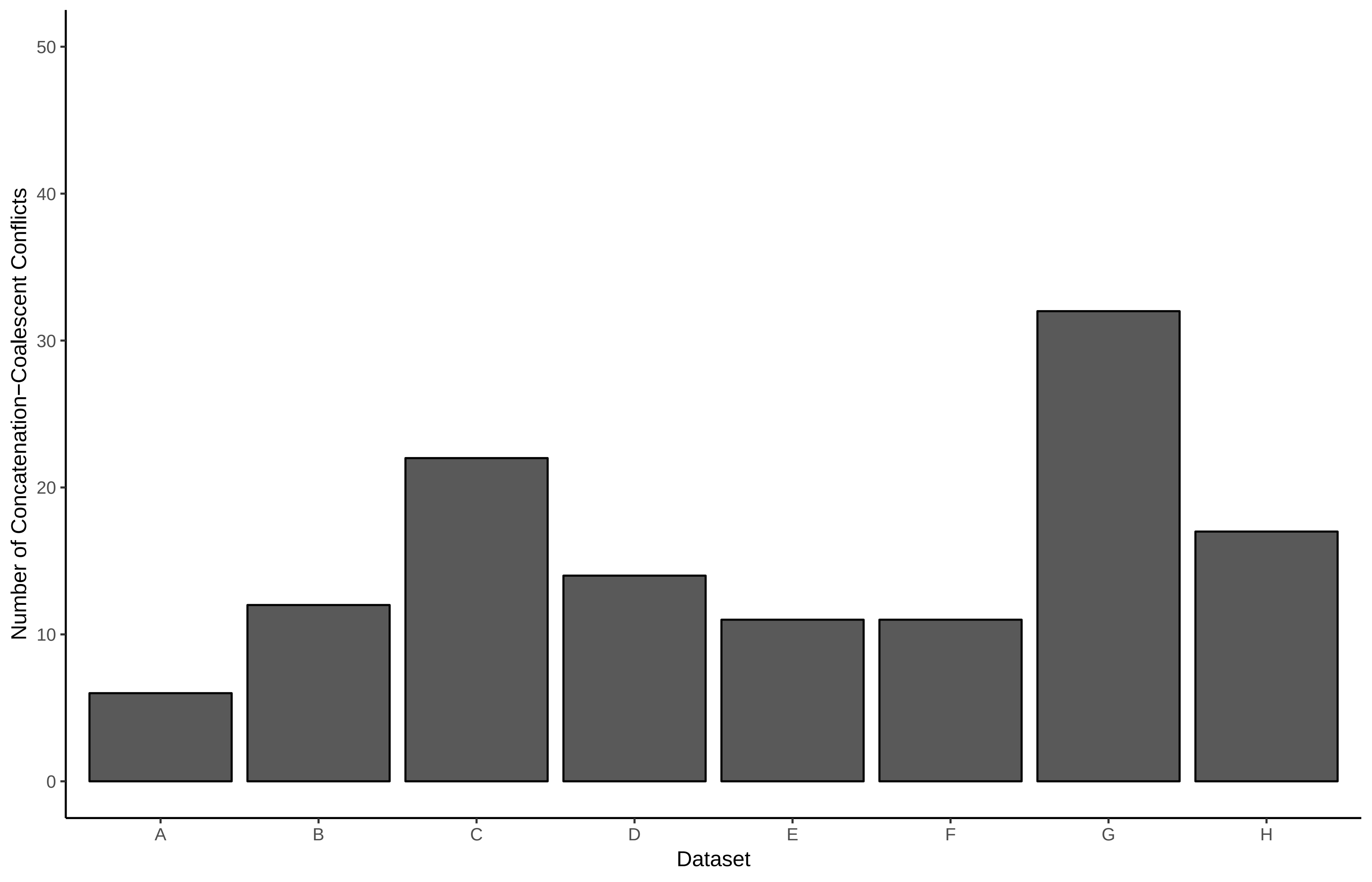

### Supplemental Figure 3

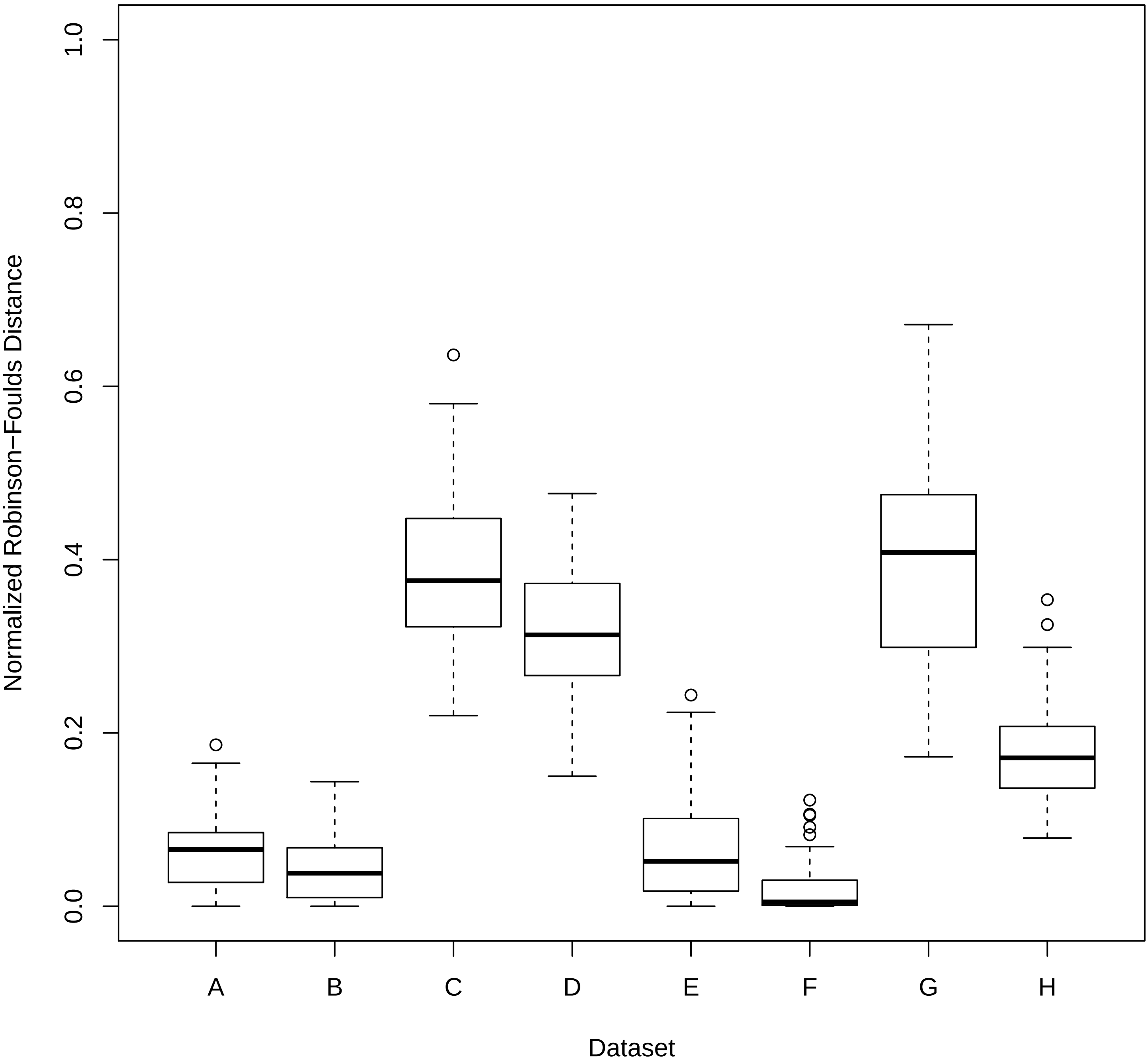

### Supplemental Figure 4

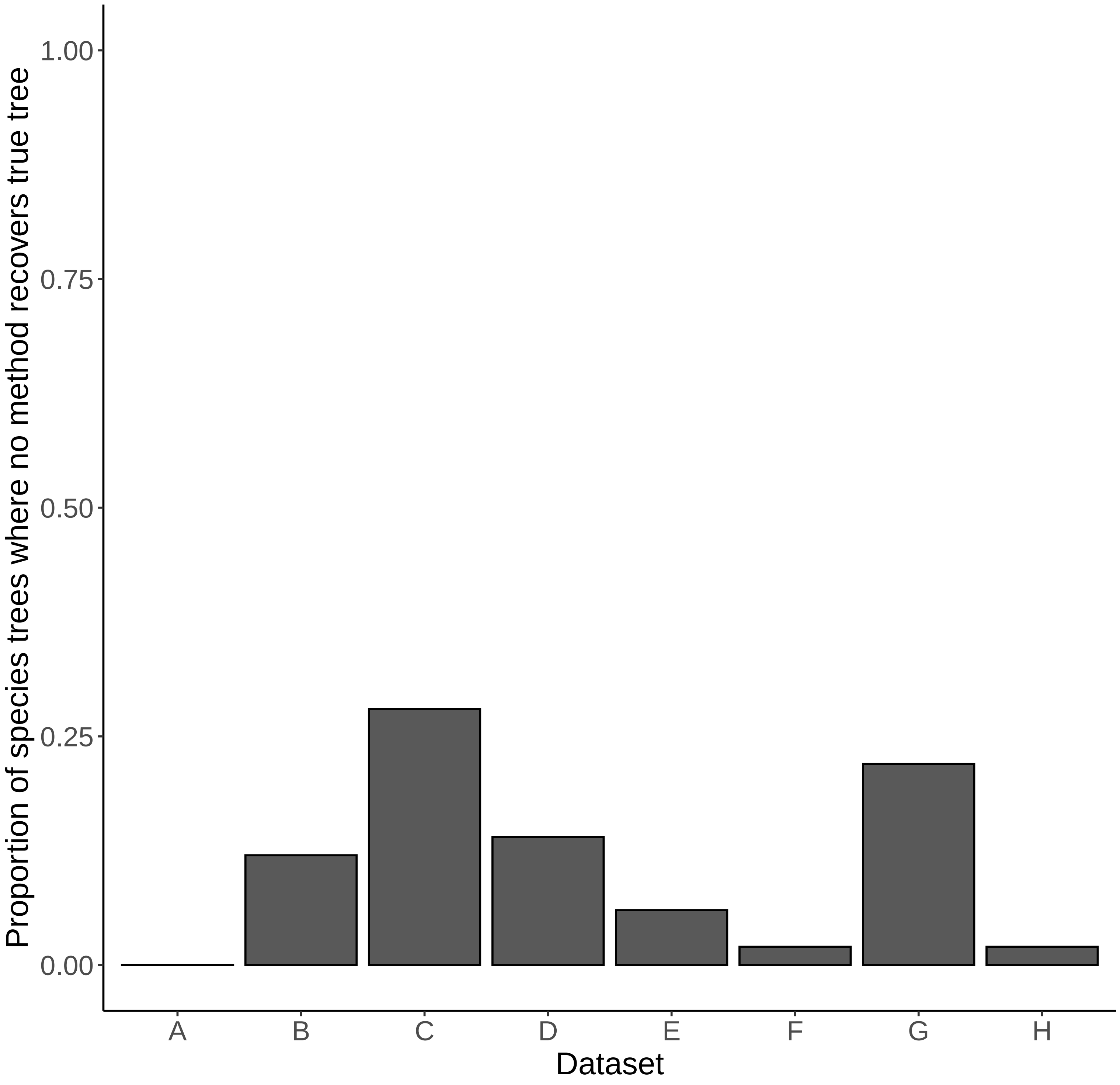

### Supplemental Figure 5

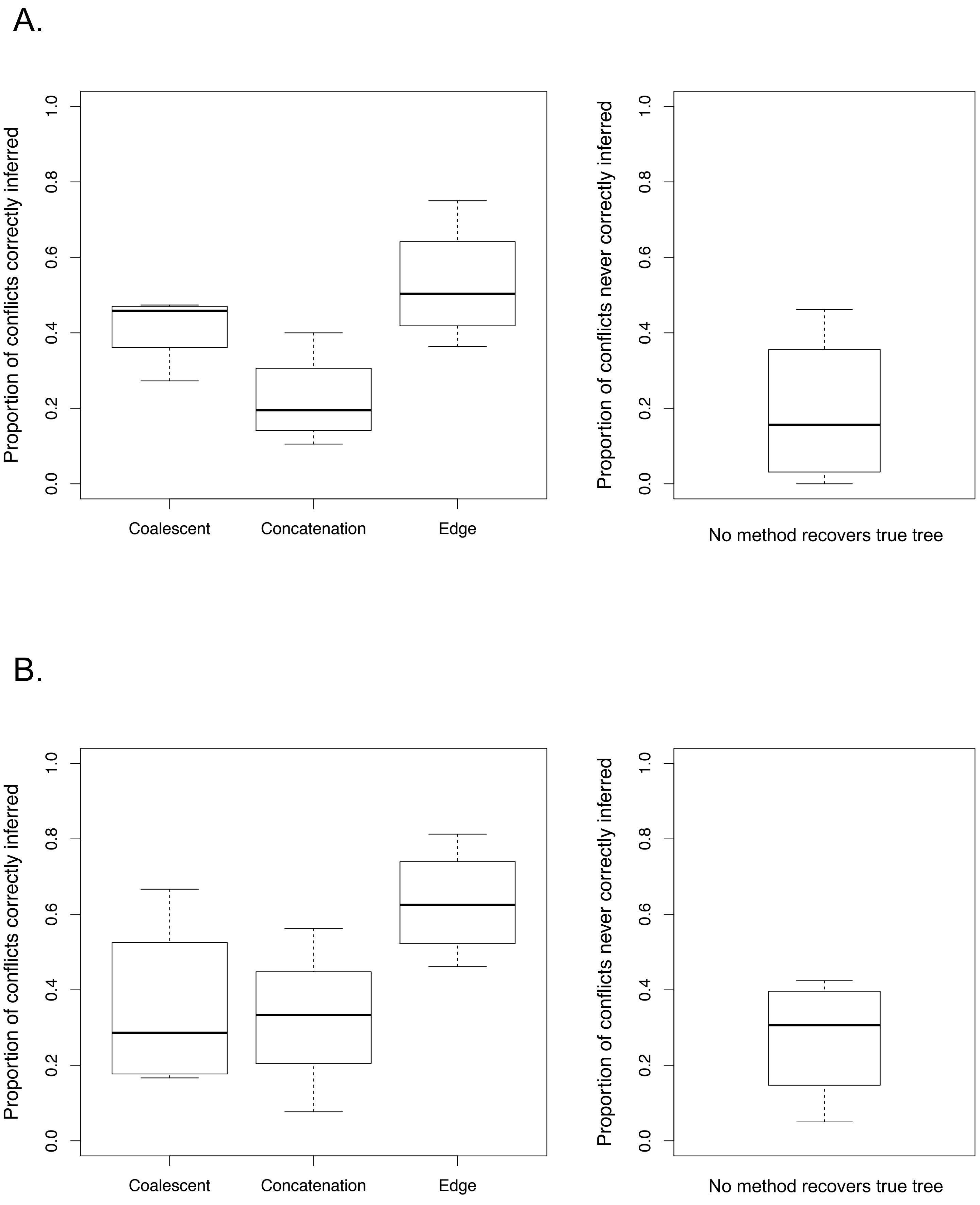

### Supplemental Figure 6

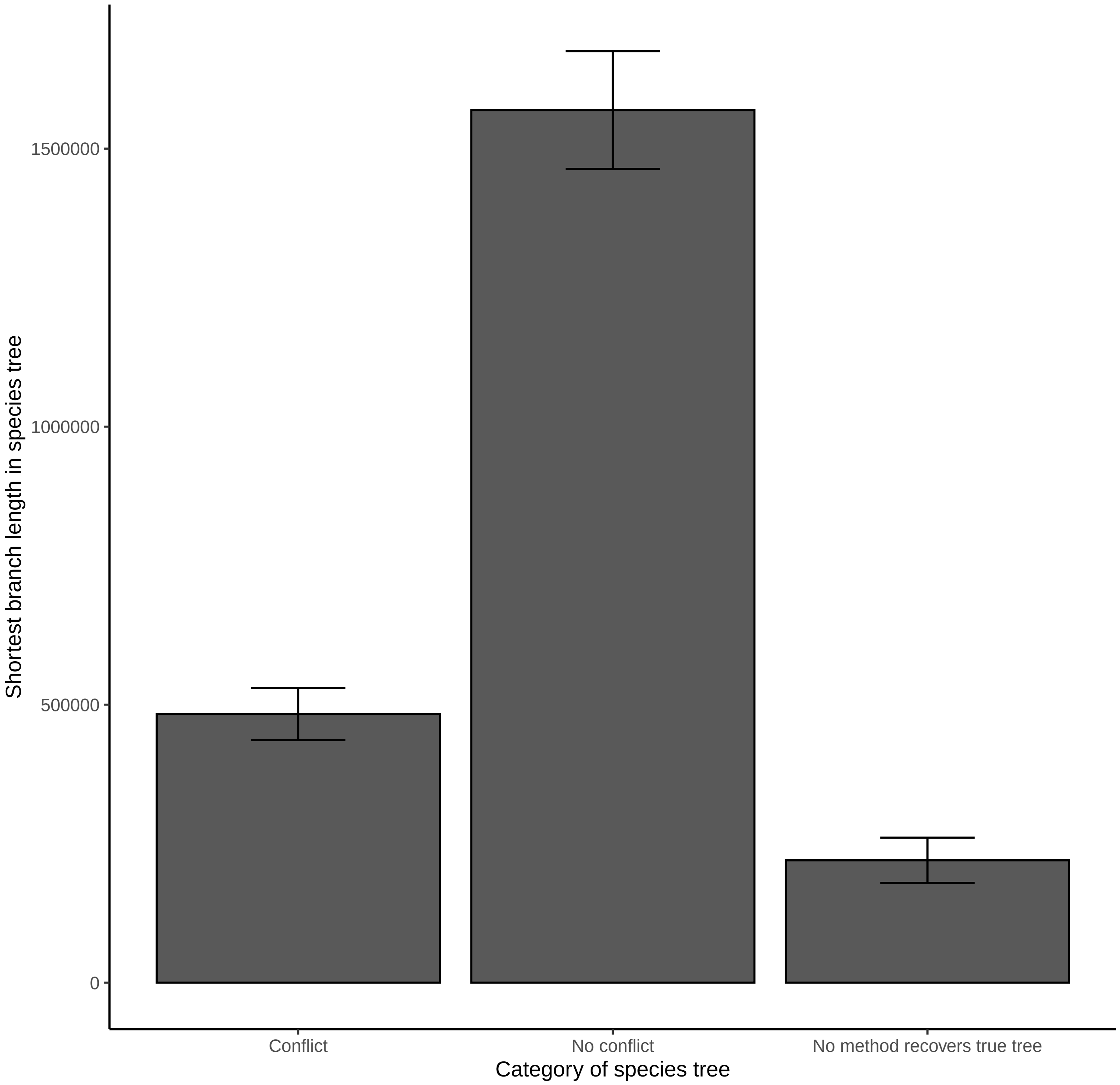

### Supplemental Figure 7

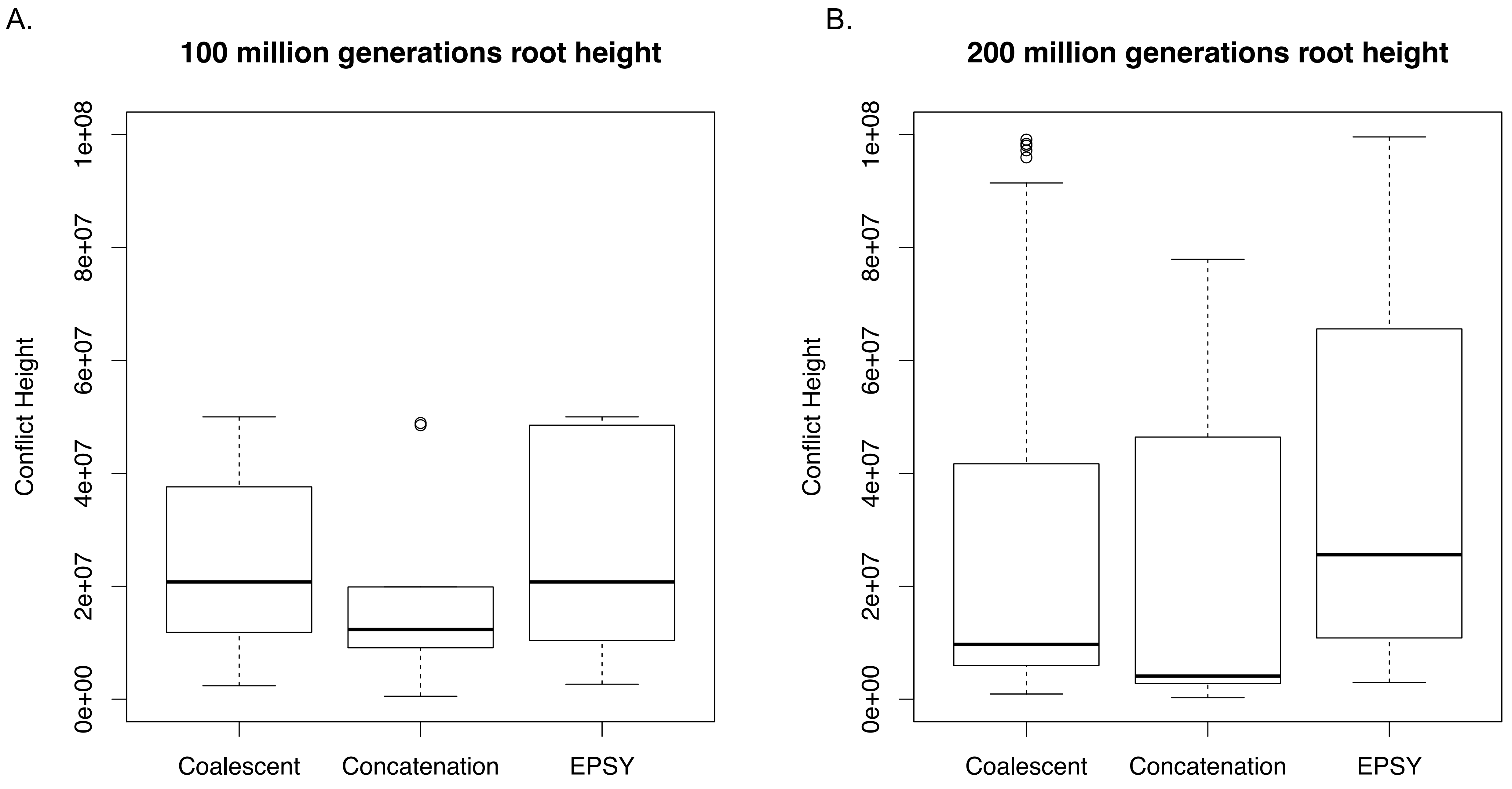
